# Structural basis of GAIN domain autoproteolysis and cleavage-resistance in the adhesion G-protein coupled receptors

**DOI:** 10.1101/2023.03.12.532270

**Authors:** Fabian Pohl, Florian Seufert, Yin Kwan Chung, Daniela Volke, Ralf Hoffmann, Torsten Schöneberg, Tobias Langenhan, Peter W. Hildebrand, Norbert Sträter

## Abstract

The GAIN domain is a hallmark of adhesion G-protein coupled receptors (aGPCRs) as this extracellular domain contains an integral agonistic sequence (*Stachel*) for activation via binding to the 7-transmembrane helical (7TM) domain of the receptor. Many aGPCRs are autoproteolytically cleaved at the GPCR proteolysis site (GPS) site within the GAIN domain formed HXS/T sequence motif. However, other aGPCR can be activated without GPS cleavage. We determined the crystal structure of the human ADGRB2/BAI2 hormone receptor (HormR) and GPCR autoproteolysis-inducing (GAIN) domains and found that this aGPCR is resistant to autoproteolysis despite the presence of a canonical HLS sequence motif at the GPS. We used structural comparisons and molecular dynamics (MD) simulations to identify structural determinants that are important for autocleavage beyond the canonical HXS/T motif. These studies characterized a conserved glycine residue and an edge-π interaction of the histidine base of the GPS sequence with a phenylalanine residue that is highly conserved in cleavage-competent aGPCRs. The MD simulations showed that this interaction is important to position the imidazole group of the histidine for deprotonation of the serine or threonine nucleophile. Removal of this interaction reduced autoproteolytic activity in the ADGRL1 receptor and restored cleavage competence of the ADGRB3 receptor in a R866H/L821F double mutant. Conservation analysis indicates that wild-type ADGRB2 and ADGRB3 are auto-cleavage-incompetent receptors.

## Introduction

The GPCR autoproteolysis-inducing (GAIN) domain is a unique feature of adhesion G-protein coupled receptors (aGPCRs), that distinguishes them from all other GPCRs ^1^. The GAIN domain bears a tethered agonist, referred to as the *Stachel*, which when released from the GAIN domain can bind to the orthosteric binding site within the 7-transmembrane helical (7TM) domain, common to all GPCRs, leading to receptor activation^2,3^. Crystal structures of the GAIN domains of human ADGRB3/BAI3 (B3) and rat ADGRL1/LPHN1 (L1) revealed a fold comprising an α-helical subdomain A followed by the C-terminal subdomain B, which consists of a twisted β-sandwich including eleven β-strands with additional short α-helices and β-strands located within loop regions^4^. Additional crystal structures of murine ADGRG1/GPR56 (G1)^5^, zebrafish ADGRG6/GPR126 (G6)^6^, human ADGRG3/GPR97 (G3)^7^ and ADGRF1/GPR110 (F1)^8^ confirmed the overall architecture of the GAIN domain fold. Proteolytic cleavage of aGPCRs was first indicated for human ADGRE5/CD97, which is processed into two noncovalently associated fragments^9^. Later, post-translational processing of murine ADGRL1/CIRL^10^ and ADGRE4/EMR4^11^ was demonstrated. The GAIN domain is sufficient for autoproteolytic cleavage of the peptide bond before the S/T residue of a canonical H-X-S/T sequence motif^4^ following a conserved mechanism^12^. The polypeptides generated by autoproteolysis at this GPCR proteolysis site (GPS) are designated as N-terminal and C-terminal fragments (NTF and CTF)^13^. The agonistic *Stachel* sequence follows directly C-terminal of the GPS, with the serine or threonine as the first residue of this sequence. CTF constructs are typically constitutively active and short synthetic *Stachel* peptides activate CTF constructs that lack the *Stachel*, resulting in an activation model involving release of the *Stachel* peptide from the GAIN domain and binding to the 7TM domain^2,3,14,15^. Notably, the *Stachel* forms a β-strand within the β-sandwich of GAIN subdomain B and appears to be tightly bound by the β-sheet and hydrophobic side-chain interactions, which stabilizes the NTF-CTF heterodimer after cleavage^4,9,11^ even when mounted on the plasma membrane^16^. The structural basis of the tethered agonist activation model was recently complemented by cryo-EM structures of the CTF of ADGRD1/GPR126^17^, ADGRF1/GPR110^17^, ADGRG2/GPR64^18^, ADGRG3/GPR97^19^, ADGRG4/GPR112^18,20^, ADGRG1/GPR56^21^, ADGRD1/GPR133^22^, ADGRG5/GPR114^22^, and ADGRL3/LPHN3^21,23^. These structures revealed a conserved binding mode of the *Stachel* sequence to the 7TM domain.

However, not all aGPCRs undergo autoproteolytic cleavage and ten of the 32 human aGPCR lack the HXS/T motif. ADGRG5 contains an HLT motif a the GPS, is cleavage-incompetent upon expression in COS-7 cells, but exhibits *Stachel*-mediated GPCR signaling^24^. While the GAIN domain structures of L1^4^, G1^5^, G3^7^ and G6^6^ were determined in the cleaved state, B3 was obtained in an uncleaved state^4^. However, B3 contains an RLS motif at the GPS and an arginine side chain should not be able to act as a base and deprotonate the serine nucleophile for autoproteolysis^12^ (Fig. 1). Based on the close homology of B2, which contains a canonical HLS motif at the GPS, to B3 we became interested in studying its structure and cleavage state to better understand the structural determinants and mechanism of GPS cleavage.

**Figure 1:**
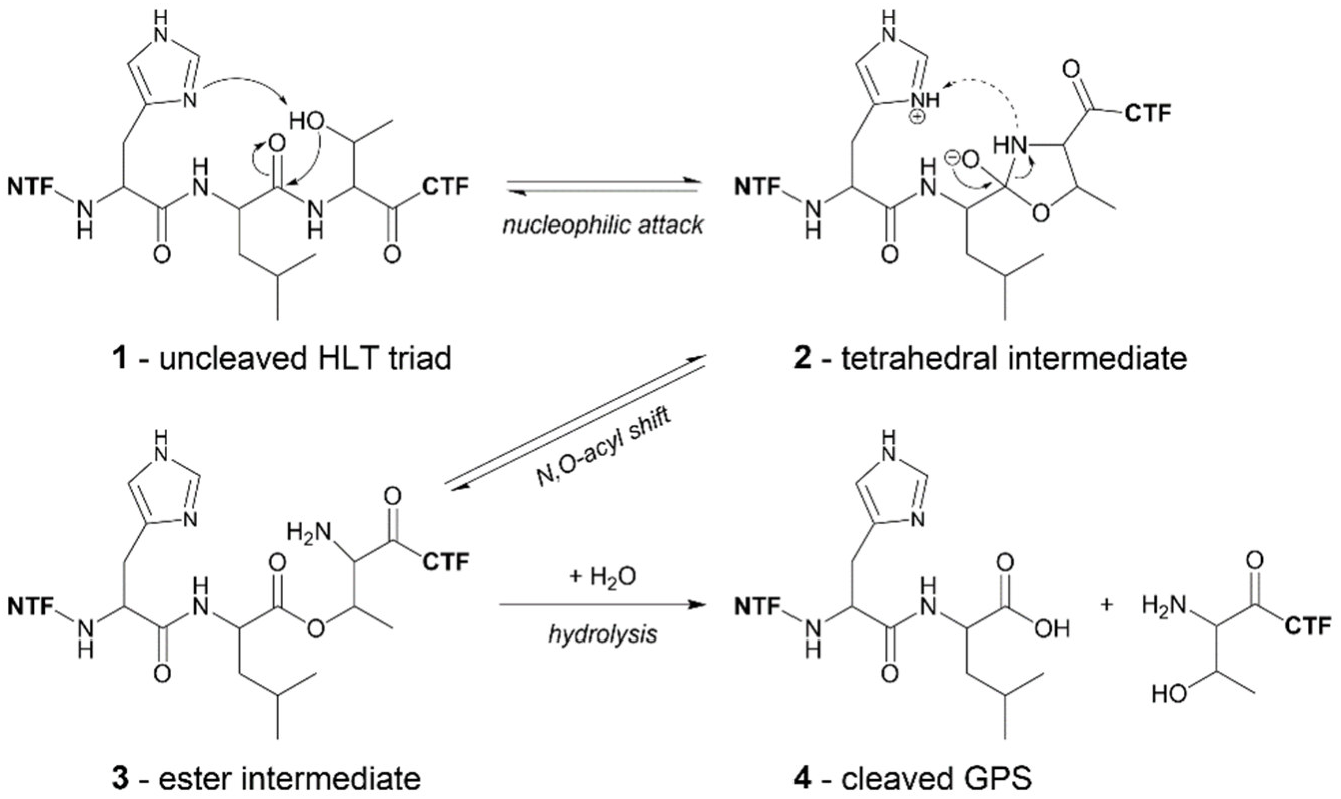
Model of the catalytic mechanism of GPS autoproteolysis with an HLT motif^12^. As outlined in the results part, a direct protonation of the nitrogen atom of the tetrahedral intermediate or transition state **2** by the histidine (dashed arrow) is unlikely due to the conformation of this structure.

Interestingly, in the crystal the B2 GAIN domain was only obtained in an uncleaved state. Molecular dynamics (MD) simulations were used to study the ensemble of conformations accessible to the GPS catalytic residues in comparison to a model of the uncleaved L1 GAIN domain. By comparing the MD structures of these two receptors and models of cleavage-competent and cleavage-resistant receptors, an edge-π interaction of the histidine base with a highly conserved phenylalanine side chain was identified, that orients the histidine base to accept a proton from the serine/threonine nucleophile. The importance of this interaction was confirmed by mutational studies.

## Results

### Expression and characterization of a full-length *hB2* ECR construct shows proteolysis at a furin cleavage site

The predicted structure of the extracellular region (ECR) of *hB2* consists in N→C-terminal order of a CUB domain, four thrombospondin (TSP) domains, a hormone receptor (HormR) domain, and a GAIN domain (Fig. 2A). No folded domains are predicted for the intracellular region of the receptor (Fig. S1A). Stable interdomain interactions are predicted to exist only between the HormR and GAIN domains (Fig. S1A). We purified and studied four constructs of the *h*B2 ECR after expression in HEK293S GnTI^-^ cells (Fig. S1B): the full-length ECR (*h*B2-ECR, corresponding to all domains <u>C</u>UB-<u>T</u>SP-<u>H</u>ormR-<u>G</u>AIN: CTHG), the ECR omitting the CUB domain (*h*B2-THG), the HormR-GAIN domains (*h*B2-HG) and the N-terminal CUB domain (*h*B2-C; Fig. S8A). For *h*B2-ECR, two fragments were observed in western blots against the C-terminal twin Strep-tag (TST), which correspond in size to proteolysis at a predicted furin cleavage site between the CUB domain and the first TSP domain (Fig. 2B).

**Figure 2:**
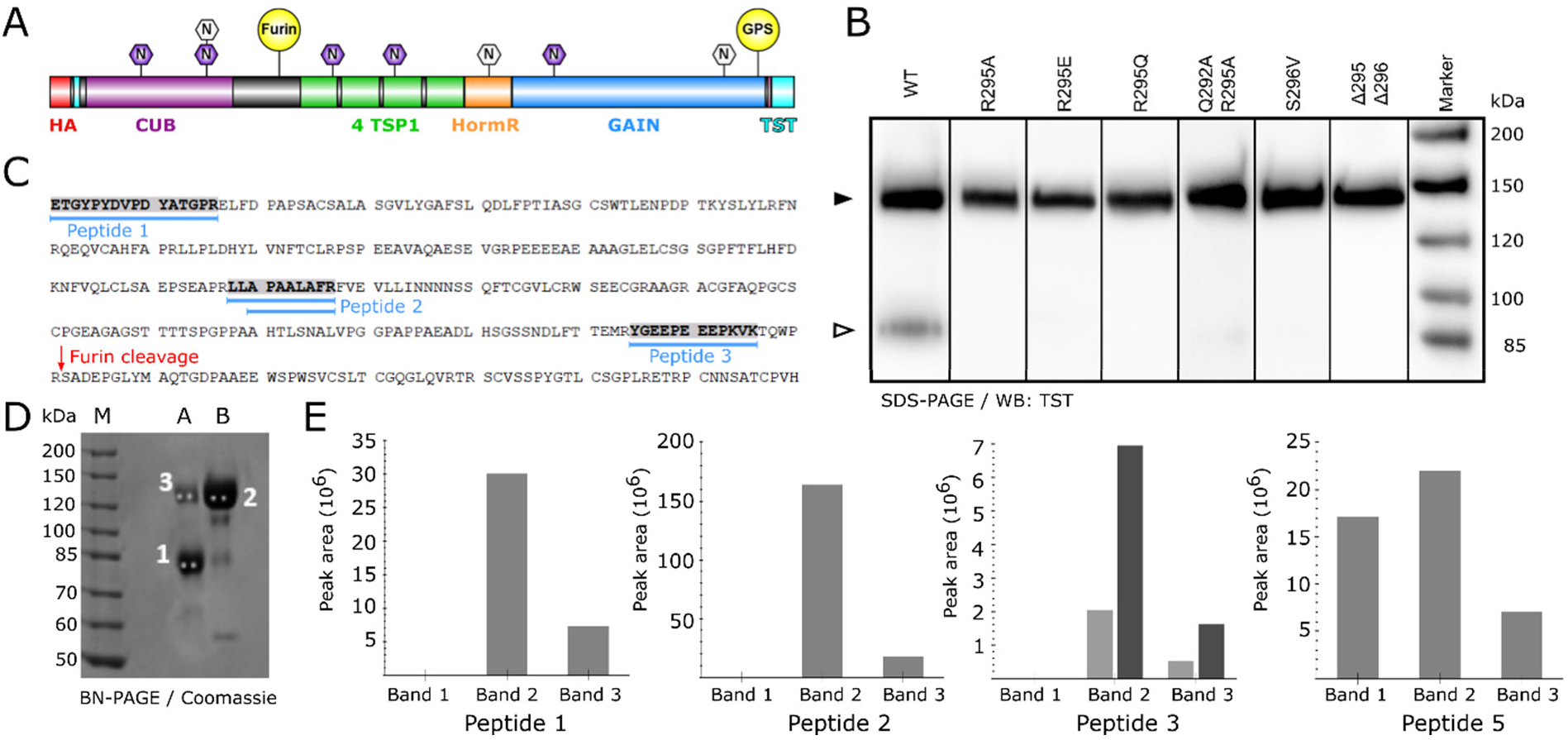
Furin cleavage of *h*B2. (A) Scheme of construct *h*B2-ECR used for these experiments. The predicted N-glycosylation sites are marked, purple marks indicate higher probability. (B) Small-scale expression tests of mutants of the furin cleavage site (Q^292^-W-P-R^296^↓) in comparison to the wild-type receptor for construct *h*B2-ECR (closed triangle, predicted mass 108.6 kDa including 8 GnTI^-^-type high-mannose N-glycosylation sites of 1.217 kDa each). The secreted proteins were detected by a western blot against the TST after a reducing SDS-PAGE. The calculated mass of the C-terminal fragment of furin cleavage (open triangle) is 78.5 kDa for this construct. Both bands appear at higher molecular masses due to extensive O-glycosylation. (C-E) Mass spectrometric identification of peptides in three bands in lanes A and B observed in a BN-PAGE of the fractions of the maxima of the first (B) and second (A) peak of a SEC chromatogram of *h*B2-ECR (similar to Figure S3). (C) Sequence of the N-terminal region of construct *h*B2. The peptides found in this region prior to the furin cleavage site are marked. (D) 2 gels plugs were cut from each Coomassie stained band and in-gel digested with trypsin followed by LC-MS/MS analysis. In total 3 peptides from position 1 to 296 and 20 peptides from position 297 to 943 of the protein were obtained. (E) Peak areas (total ion current for the mass range selected for the specific peptide) detected for peptides 1-3 and peptide 5 (of the region after the furin cleavage site, as control, Fig. S2) for each of the three bands. For peptide 3 two signals were observed due to different charges.

Furin is a proprotein convertase expressed in a broad range of mammalian cell types, responsible for the conversion of peptide hormones, neuropeptides and other proteins into their biologically active form^25^. Although the furin recognition site of *h*B2 (Q^292^-W-P-R^296^ ↓) deviates from the canonical recognition site (R-X-K/R-R ↓)^25,26^ partial cleavage was observed in small-scale expression tests of the wild type construct (*h*B2-ECR) and later during large-scale expression and purification. To abrogate furin cleavage, various point mutations were introduced in the recognition site and small-scale expression tests verified their efficacy (Fig. 2B). The double mutant (Q292A/R295A; *h*B2-ECR-DM) was chosen for large-scale transient expression and purification by affinity chromatography via the twin Strep-Tag. The amount of cleaved and uncleaved protein was estimated from size-exclusion chromatography (Fig. S3). Approximately 15.6 % of the wild-type protein was cleaved by furin, while 7.4 % were cleaved of the double mutant, following transient expression and purification. Expression of the wild-type protein in a stable cell line led to an increase of the furin-cleaved fraction to approximately 28.8 % (data not shown).

### All B2 ECR constructs form homodimers and are not cleaved at the GPS

We purified all four constructs to homogeneity by Strep-Tactin affinity chromatography followed by size exclusion chromatography (SEC) (Figs. S4-S8). In blue-native polyacrylamide gel electrophoresis (BN-PAGE) analysis, all constructs of *h*B2 were observed at sizes too large for just a monomer (Fig. S9, **Table** 1). In the case of *h*B2-ECR, bands at 293 kDa and 188 kDa indicated the presence of two homodimers, one of the full-length protein and one of the C-terminal fragment following furin cleavage. Interestingly, a band corresponding to a heterodimer of one cleaved and one uncleaved chain was not observed indicating that processing by the furin protease always affects both chains or none. Furthermore, no exchange between furin-cleaved and unprocessed subunits occurs after secretion. This is demonstrated by (*i*) mass spectrometry analysis of three peptides in the N-terminal region of the cleavage site (Fig. 2), (*ii*) comparison of the band intensities of a western blot of the TST with the SEC elution profile (Fig. S4) and (*iii*) a western blot against the HA-Tag (Fig. S10). A homodimer was also observed for *h*B2-THG at a size of 188.8 kDa. Bands of *h*B2-HG were very broad, but centered around 96.5 kDa, again close to the expected size for a homodimer. Although not as clear-cut as for the other constructs, *h*B2-C, observed at 83.3 kDa, also appeared too large for just a monomer. Estimation of the particle size by dynamic light scattering (DLS) supported the finding that constructs of *h*B2-ECR, *h*B2-THG and *h*B2-HG form homodimers in solution (Fig. S7, Table S1).

**Table 1:** Comparison of estimated and observed molecular weights of *h*B2 constructs. Monomer sizes were determined by SDS-PAGE under reducing conditions and putative dimer sizes were observed in BN-PAGE. ^1^denotes the full-length ECR; ^2^denotes the C-terminal fragment following furin cleavage. The calculated molecular weight is based on the mass of the amino acids and the putative N-glycosylation sites (1.217 kDa per high-mannose N-glycosylation site). O-glycosylation may contribute to the deviations between observed and calculated molecular masses. ^3^The two values denote the sizes of a *h*B2-ECR after furin cleavage of one or two chains of the *h*B2-ECR dimer, respectively.

| Construct name | Construct residues | Monomer mass [kDa] (calculated) | Monomer mass [kDa] (observed, SDS-PAGE) | Dimer mass [kDa] (calculated) | Dimer mass [kDa] (observed, BN-PAGE) |
| --- | --- | --- | --- | --- | --- |
| *hB2-ECR* | F33-P921 | 108.6<sup>1</sup> | 134.9<sup>1</sup> | 217.2<sup>1</sup> | 293.0<sup>1</sup> |
|  |  | 78.5<sup>2</sup> | 87.4<sup>2</sup> | 180.5<sup>3</sup>, 143.8<sup>3</sup> | 188.0<sup>2</sup> |
| *hB2-THG* | S296-P921 | 79.7 | 87.3 | 159.4 | 188.8 |
| *hB2-HG* | P528-P921 | 55.4 | 52.3 | 110.8 | 96.5 |
| *hB2-C* | F33-R295 | 36.7 | 52.0 | 73.4 | 83.3 |

The western blots against the C-terminal TST (Fig. 2, S4-S6) or the N-terminal HA-tag (Fig. S10) after SDS-PAGE demonstrated intact GPS without autoproteolysis for all constructs containing the GAIN domain.

### Crystal structure of the B2 HormR and GAIN domains

The purified construct *h*B2-HG exhibited favorable monodispersity and a high melting temperature of 76.6±0.08°C (Fig. S7). It could be crystallized (Table S2) and the crystal structure was determined to 2.2 Å resolution (Figs. 3 and S11, Table S3). The model comprises residues 532 to 921. Two loops of residues 605-609 and 770-810 are disordered. In agreement with the gel electrophoresis experiments, the electron density showed that no cleavage has occurred at the GPS. The GPS is located at the bottom of a groove, which is flanked by residues 848 to 856 (flap 1) on one side and 879 to 886 (flap 2) on the other side of the groove (Figs. 3B, S13). These flaps have been shown to be flexible in molecular dynamics (MD) simulations of ADGRGL1, ADGRG1 and ADGRE5 resulting in open states, in which the GPS is well accessible from the solvent, and closed states, in which in particular flap 1 covers the GPS^16^. Notably, the long disordered loop region 770-810 of B2 is located next to flap 1 and may contribute to its flexibility. The *h*B2-HG crystal structure represents an open state concerning the flap conformations (Fig. 3B). A cacodylate ion from the crystallization buffer has bound in the groove next to the GPS. H910 of the GPS HLS sequence motif is present in two alternative conformations (Fig. S12). In the main conformation, which is shown in Figure 3C, the imidazole group is oriented closer to the nucleophilic S912 side chain. For a direct hydrogen bonding interaction between S912 and H910, which is required for proton transfer as the assumed first step of GPS hydrolysis (Fig. 1), S912 needs to change conformation from the t-rotamer observed in the crystal structure to the m-rotamer and the imidazole group of H910 needs to flip. The side-chain flip of H910 in the crystal structure was modeled based on a hydrogen bonding interaction of the N_ε_-atom with a nearby water molecule. Thus, neither the S912 nucleophile nor the H912 base are positioned to initiate the cleavage reaction in the crystal structure. However, both residues are expected to be able to change conformation as there are no steric clashes preventing these conformational changes. If the rotamers are manually changed to the energy minimum of a serine m-rotamer (χ_1_ = −65.0°) and to a flipped H910 side chain, the H910 N_δ_-atom and the S912 O_γ_ are 2.8 Å apart and positioned for a hydrogen bonding interaction.

**Figure 3:**
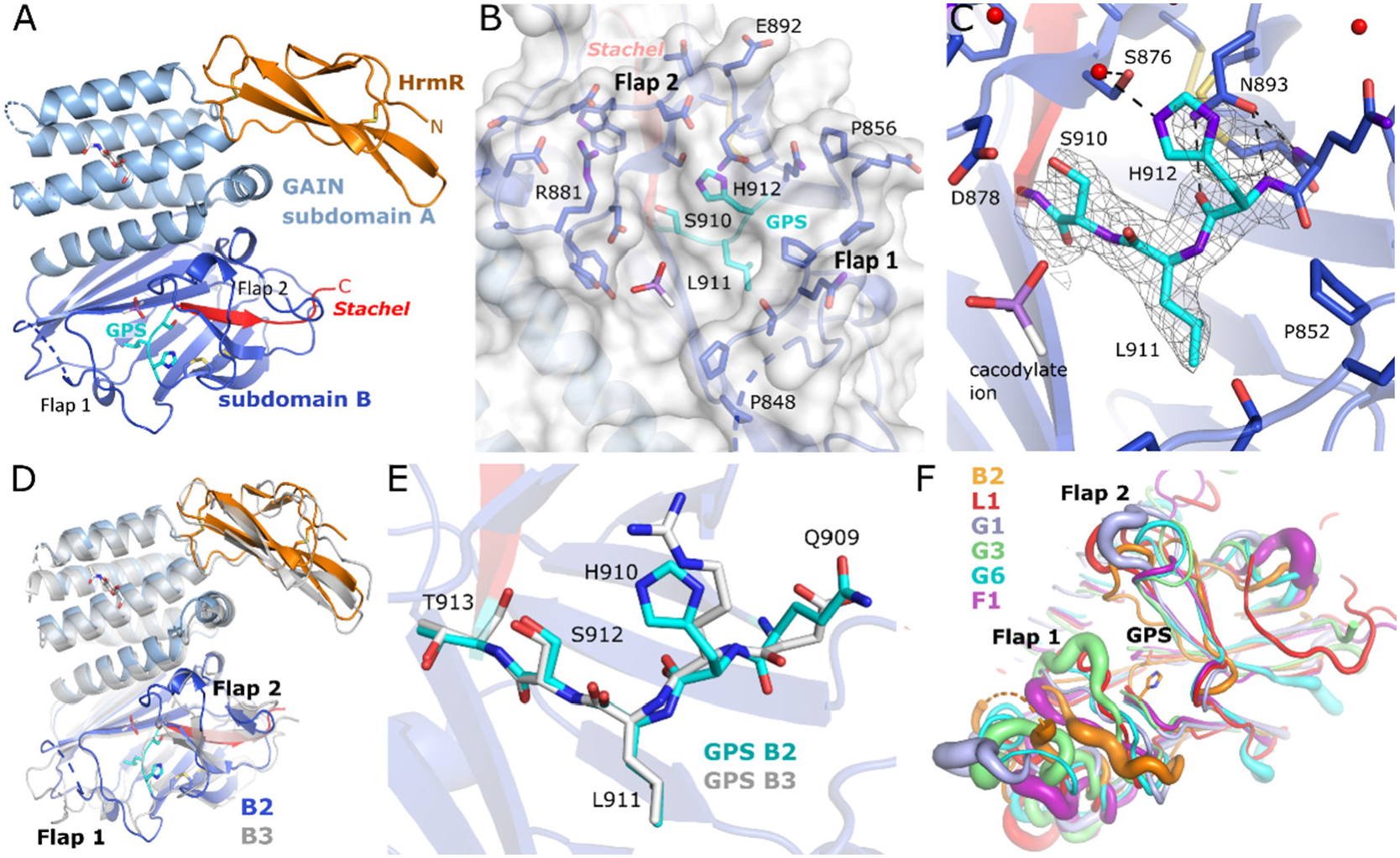
Crystal structure of the B2 HormR and GAIN domains and structural comparisons. (A) Overview of the domain structures. One disulfide bridge is located in the HormR domain and two cystine linkages are found in the GAIN subdomain B. (B) The GPS is solvent accessible at the bottom of a groove. A cacodylate ion from the crystallization buffer binds in the groove. (C) Close-up view of the GPS. The (2F_o_-F_c_)-type electron density of the GPS residues is shown at a contour level of 0.8. For H912 only the major conformation is shown. (D) Comparison of the crystal structures of the HormR and GAIN domains of B2 (in color) and B3 (grey) (E) Comparison of the GPS structures of B2 and B3 (PDB id 4dlo^4^). The GAIN domain structures have been superimposed based on subdomain B. (F) Superposition of the crystal structures of B2, L1, G1, G3, and G6. The thickness of the tube indicates the magnitude of the crystallographic B-factors, which increase mostly with conformational flexibility in the crystal. The view focuses on the GPS and the flanking flaps. All superimposed structures were aligned to the GAIN subdomain B.

A comparison of the HormR-GAIN domain structures of B2 and B3 shows a similar relative orientation of the two domains (Fig. 3D). The disordered loop of subdomain B is much shorter in the B3 structure and only two residues were not modeled due to flexibility. There are no significant differences in the overall fold of the two related GAIN domains, which share 49.4 % sequence identity and superimpose with an rmsd of 2.22 Å for all common 340 C_α_-atoms and 1.38 Å for 298 C_α_ atoms (omitting some deviating loop conformations). The secondary structure elements superimpose fairly well, but interestingly the largest differences in the main chain structures are observed in the regions surrounding the GPS (Fig. 3D). While the main chain structures of the GPS of B2 and B3 match closely (Fig. 3E), the immediate environment differs in many residues and interactions (Fig. S13). A comparison to the crystal structures of cleaved GAIN domains shows that flap 1 is closed in L1, G1, G3, and G6 (Fig. 3F).

This comparison also demonstrates the conformational diversity of the two flaps and the loop next to flap 1. In addition, these regions also often display increased B-values resulting from a limited flexibility in the crystal (Fig. 3F). It has been noted for the B3 GAIN structure that threonine at the GPS +1 position (the position directly after the cleavage site) is an outlier in the Ramachandran plot in both chains observed in this crystal structure^4^. This situation of a strained main chain conformation at S912 is also observed in the B2 GAIN domain with main chain torsion angles of ψ = −159.5° and φ = −179.1°.

**The GAIN domain of ADGRB2 is resistant to GPS cleavage even at elevated temperature or in the presence of hydroxylamine**

Following expression in HEK293S GnTI^-^ cells, all constructs of B2 containing the GAIN domain were always detectable via their C-terminal TST and no traces of cleaved NTFs were detected throughout the purification process. As the crystal structure analysis demonstrated a properly folded GAIN domain and GPS structure, we tested if thermal activation might be efficient to cross an activation barrier of the autoproteolysis reaction. Incubation of the protein up to one hour at 60°C did not result in GPS cleavage (Fig. S14). For other systems of *cis*-autocleavage, it has been observed that the reaction can be stalled at the ester intermediate state^27^. Although an ester intermediate was not observed in the crystal structure, there might be an equilibrium between the highly populated initial peptide ground state and a higher energy ester intermediate, which cannot proceed via the final hydrolysis step. Such ester intermediate states can be hydrolyzed by added hydroxylamine^27^. We added up to 500 mM hydroxylamine to the purified *h*B2-HG construct, however no GPS cleavage was observed (Fig. S14).

### Analysis of possible factors influencing GAIN domain autoproteolytic activity

To understand the structural determinants that might contribute to the autoproteolysis resistance of the B2 GAIN domain, we compared it to cleavage-competent aGPCRs. Because structures of pre-cleavage states of autoproteolytically active GAIN domains are not available, we resorted to a comparison to AlphaFold models. We generated models for the GAIN domains of receptors G1, G3, G6 and L1, for which crystal structures of GAIN domains in the cleaved state are available and for which the cleavage competence has been demonstrated for the full-length receptors of G1^28^, G6^29^ and L1^4,10^.

A comparison of the *h*B2-HG crystallographic and AlphaFold models with the AlphaFold models of the precleavage states of B2, L1, G1, G3, and G6 indicates minor differences in the conformation of the main chain of the GPS region in addition to the different conformations of the flap regions (Fig. 4A). A striking difference in the immediate environment of the GPS concerns residues S876 and N893 as well as the main chain fold of the region around N893 (Fig. 4B). S876 is replaced by a phenylalanine in the autoproteolysis-competent receptors compared in Figure 4B, but also in other receptors, for which GPS-autoproteolytic activity has been detected in previous work (Fig. S15). The side chain of this phenylalanine (or tyrosine in E3) interacts with the histidine base of the GPS in a T-shaped π-π interaction. N893 is unique in B2 and it interacts with the H910 side chain via π-stacking and with hydrogen bonds to the main chain atoms (Fig. 3C). This residue corresponds to a strictly conserved glycine in the autoproteolysis-competent human receptors and the glycine is part of a region near the GPS that shows little differences in the AlphaFold models (Fig. 4B and S15A). The next residue is C894, which is part of a disulfide bridge that is present in all human aGPCR GAIN domains.

**Figure 4:**
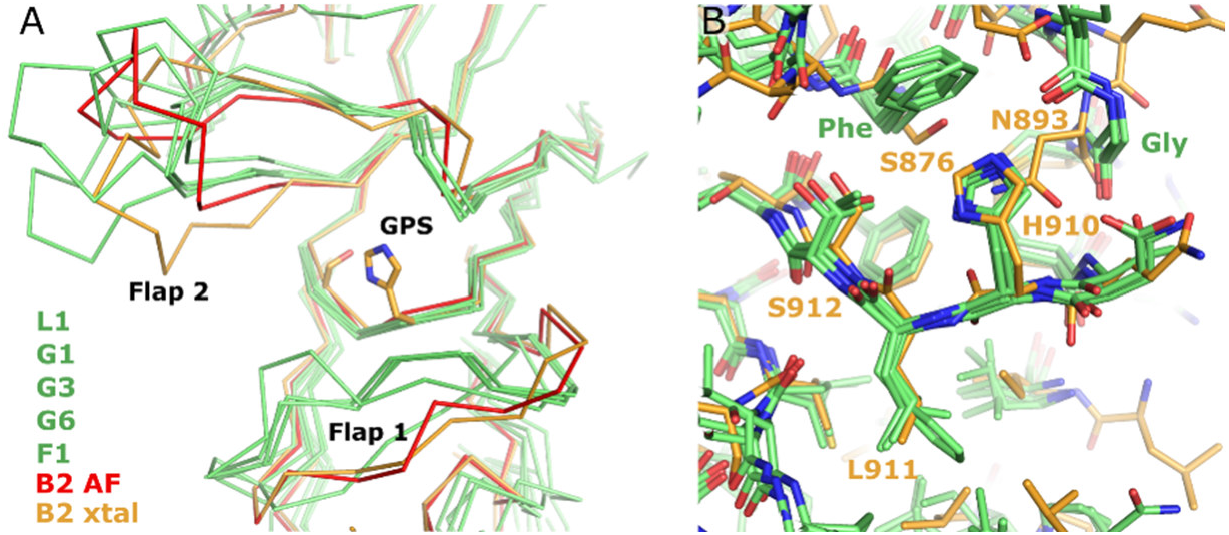
Comparison of the GPS and its environment between the B2 GAIN domain and cleavage-competent receptors. (A) Superposition of the C_α_-traces of the B2 GAIN domain crystal structure (orange) with its AlphaFold model (red) and the AlphaFold models of the uncleaved GAIN domains of L1, G1, G3 and G6 (all green), for which crystal structures of cleaved GAIN domains have been determined. (B) Comparison of the GPS and its environment between these GAIN domain models. The B2 AlphaFold model is omitted here.

Considering the mechanistic model of GPS autoproteolysis^12^ (Fig. 1), the following factors may contribute to the catalytic efficiency of the cleavage reaction: (*i*) the protonation state of the histidine base, (*ii*) the relative orientation of the alcohol (Thr or Ser) nucleophile and the histidine for efficient proton transfer, (*iii*) the position of the hydroxyl group for nucleophilic attack on the carbonyl carbon of the adjacent peptide bond, (*iv*) strain of the peptide bond (*i*-*iv* in the ground state **1**), (*v*) stabilization of the tetrahedral transition state or intermediate (**2**), (*vi*) protonation of the leaving amine group for efficient transfer to the ester intermediate (**3**), (*vii*) position and activation of a water nucleophile for attack on the carbonyl group of the ester intermediate, (*viii*) protonation of the alcohol leaving group of the threonine or serine side chain of the CTF (**4**) in the cleaved protein, and (*ix*) stabilization of the tetrahedral intermediate of ester hydrolysis.

In the AlphaFold models of L1, G1, G3, and G6 the histidine base was present in the p80-rotamer, where the N_δ_-nitrogen is facing towards the Ser/Thr nucleophile. Only for G6, the threonine nucleophile was predicted in the p-conformation and at a distance of 3.3 Å positioned to interact with the histidine base for proton transfer. However, similar to the situation described above for the B2 GAIN domain crystal structure, the Ser/Thr nucleophiles are free to adopt other rotamers. We therefore analyzed the geometric requirements for efficient attack of the nucleophile on the carbonyl group of the adjacent peptide bond for conformations observed in molecular dynamics (MD) simulations. The computational estimation of pK_a_ values is also strongly dependent on details of the residue environment and is therefore preferably assessed for an ensemble of available conformations observed in MD simulations.

### Molecular dynamics simulations on the ADGRB2 and ADGRL1 GAIN domains

Cleavage-competent conformations of the GAIN domain may differ from low-energy states observed in crystal structures or in energy-minimized structural models. We therefore employed molecular dynamics (MD) simulations to analyze and compare the conformational landscapes of the L1 and B2 GAIN domains for factors that may influence the initial step of GPS autoproteolysis. A model for the uncleaved rat L1 GAIN domain was generated based on the crystal structure of the cleaved L1 GAIN domain^4^. The histidine base must be in the deprotonated neutral state to act as a general base. We tested the behavior of the L1 and B2 receptor each in three protonation states HSE (proton at N_ε_), HSD (proton at N_δ_) and HSP (both nitrogens protonated) in three 1500 ns replica simulations. The simulations reached a stable conformation indicated by a plateau of backbone-RMSD values after about 100-200 ns (Fig. S16). Larger RMSD values of ~6-7 Å were observed for L1 compared to B2 (~2-3 Å). This is primarily due to much larger fluctuations of loop regions in L1 (Fig. 5). The HormR domain shows a rigid-body motion relative to the GAIN domain and a large mobility is observed in particular for flap 2. The loop between residues 605 and 609 also shows high mobility in agreement with its missing electron density in the crystal structure. An analysis of the predicted pK_a_ values of H910 in B2 and H836 in L1 shows that the most likely protonation state is HSE for both GAIN domains with average pK_a_ values of around 3 (Fig. S17).

**Figure 5:**
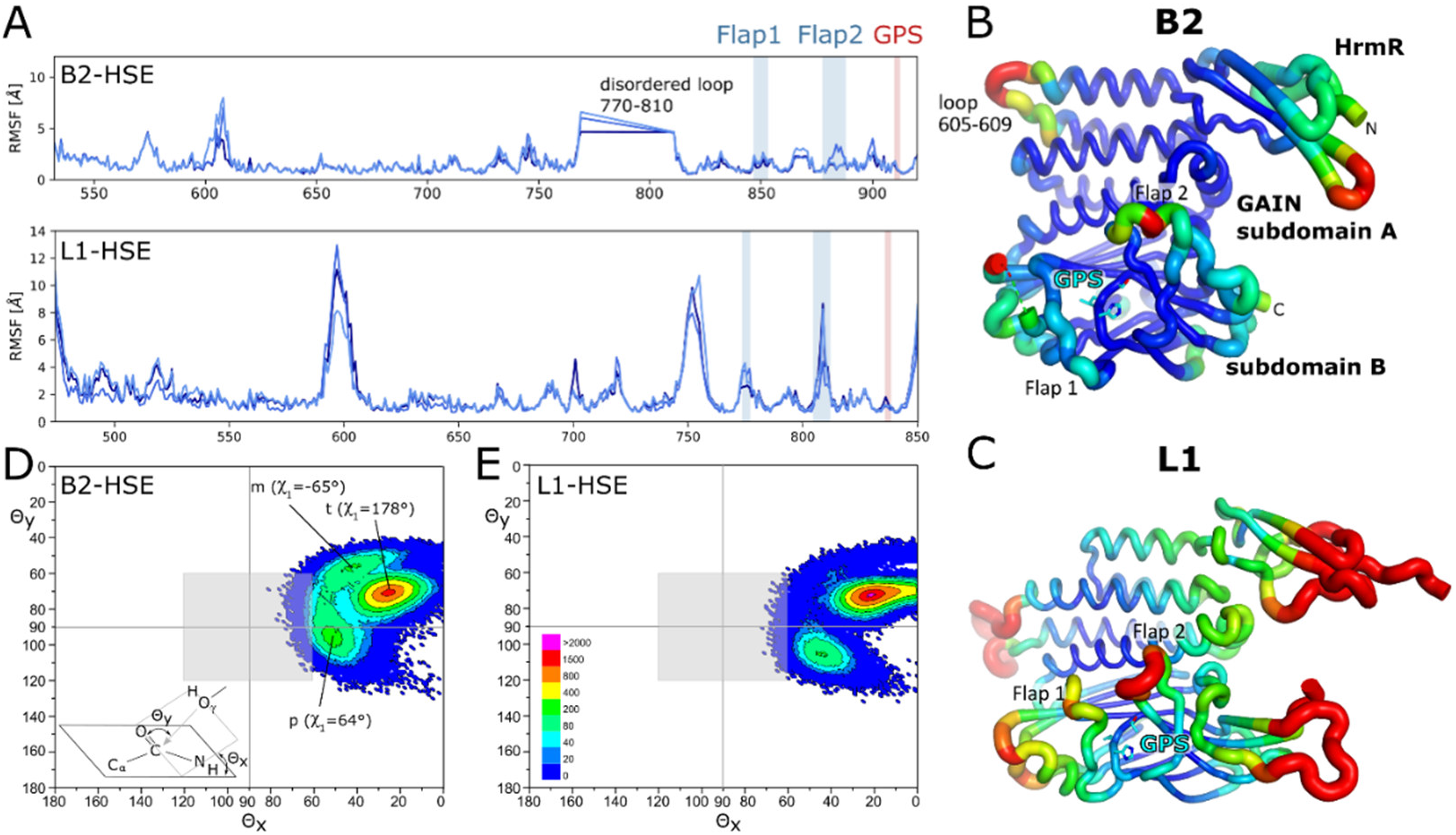
Mobility of residues in the MD simulation. (A) Plot of the root mean square fluctuations (RMSF) of the C_α_-coordinates of individual residues of B2 (top) and L1 (bottom) for three replicas of the HSE simulations (dark blue: r1, medium blue: r2, bright blue: r3). (B) RMSF values of the C_α_-coordinates mapped onto the fold of the HormR-GAIN domains such that high mobility is indicated by red color and thick tube regions and low mobility by blue color and thin tube regions. The RMSF values were calculated for all three replicas of the B2-HSE simulations. (C) RMSF values of all three replicas of the L1 HSE simulation mapped onto the fold of the L1 HormR-GAIN domains. (D) and (E) Scatterplots for the orientation of the hydroxyl nucleophile (S912 for B2 and T839 for L1) relative to the carbonyl carbon atom of the scissile peptide bond in MD simulations of the B2 and L1 GAIN domains. The definition of angles θx and θy is shown in the scheme left and was adapted from^30^. (D) Scatterplot for in total 452021 frames of the three replicas of the B2 simulations with H910 protonated at N_ε_. (E) Scatterplot for in total 452124 frames of the three replicas of the L1 simulations with H837 protonated at N_ε_. The grey square indicates the arbitrary area of θ_x_ = 90° ± 30° and θ_y_ = 90° ± 30° used to quantify the number of frames with a favorable geometry of the hydroxyl group for nucleophilic attack on the carbonyl carbon in Table 2. The scale indicates the color coding for both plots to visualize the number of frames for which a given pair of θx and θy (binned in 1° steps) is observed.

Radisky and Koshland^30^ analyzed the position of the attacking alcohol nucleophile in complexes of serine proteases with peptide or protein inhibitors, which presumably can react to form acyl-enzyme intermediates in equilibrium with the Michaelis complex but are not completely hydrolyzed due to a slow deacylation step. In 79 structures the nucleophile positions were tightly clustered at angles θ_x_ = 90° and θ_y_ = 90° perpendicular to the peptide plane at the carbonyl carbon position. This orientation compares to an optimal angle of θ_y_ = 105° derived from theoretical calculations^31^. A comparison of the scatterplots depicting the positions of the alcohol nucleophile relative to the carbonyl group for the simulations of B2 and L1 (Fig. 5D/E) shows that in both simulations the nucleophile is mostly located at θ_x_ ≈ 25° and θ_y_ ≈ 72°. In the simulation of L1, however, the nucleophile less often deviates from this position compared to B2. Consequently, for the B2 GAIN domain the nucleophile approaches more often a position favorable for nucleophilic attack (Table 2).

**Table 2:** Geometric parameters describing the orientation of the S/T alcohol nucleophile, the histidine base and the scissile peptide bond in the MD simulations of the B2 and L1 GAIN domains. The values specify the number or percentage of frames in which the given condition is satisfied. These simulations were carried out with a neutral histidine base protonated at N_ε_ (HSE). Results from simulations with protonation at N_δ_ are shown in Table S4. The underlined χ_1_ values denote rotamers, in which a hydrogen bonding interaction between the histidine and the S/T alcohol nucleophile is possible. B2FG denotes the S876F/N893G B2 variant.

| | $\Theta_x = 90^\circ \pm 30^\circ$<br>and<br>$\Theta_y = 90^\circ \pm 30^\circ$ | Hbond* | Hbond<br>and<br>$\Theta_{x/y} = 90^\circ \pm 30^\circ$ | $\chi_1^{**}$ His<br><u>$+60^\circ$</u>, $-60^\circ$, $-180^\circ$ | $\chi_1^{**}$ Ser/Thr<br><u>$+60^\circ$</u>, <u>$-60^\circ$</u>, $-180^\circ$ |
| --- | --- | --- | --- | --- | --- |
| **B2 r1** | 3597 | 19 | 1 | 1.9%, 5.2%, 69.3% | 26.2%, 19.8%, 45.6% |
| **B2 r2** | 1020 | 9 | 0 | 4.2%, 17.2%, 50.3% | 4.9%, 17.3%, 67.6% |
| **B2 r3** | 2876 | 35 | 0 | 1.1%, 14.0%, 63.6% | 21.9%, 14.1%, 55.7% |
| **L1 r1** | 147 | 1807 | 23 | 60.2%, 33.9%, 2.7% | 4.8%, 0.7%, 60.5% |
| **L1 r2** | 35 | 1351 | 3 | 98.1%, 0.7%, 0.0% | 2.6%, 1.7%, 59.4% |
| **L1 r3** | 359 | 7469 | 33 | 95.8%, 1.8%, 0.7% | 16.0%, 2.9%, 53.3% |
| **B2FG r1** | 517 | 268 | 16 | 10.2%, 53.9%, 25.2% | 4.5%, 6.3%, 64.5% |
| **B2FG r2** | 537 | 538 | 14 | 7.3%, 57.5%, 25.0% | 1.5%, 5.8%, 69.7% |
| **B2FG r3** | 51 | 1 | 0 | 1.6%, 77.8%, 14.7% | 0.1%, 3.1%, 70.8% |

\*A favorable hydrogen bonding geometry was defined as a distance of 3.2 Å or less between the histidine nitrogen atom and the Ser/Thr O<sub>γ</sub> atom and a deviation of less than 30° of the O<sub>γ</sub>-H...N angle from linearity.
\*\*The $\chi_1$ torsion angle was assigned to the given value if it deviated by less than 30° from this value.

In contrast, a hydrogen bonding interaction of the histidine base and the hydroxyl nucleophile is more frequently observed in the simulation of L1 (Table 2). As a result, the L1 receptor adopts rarely, but much more often than B2, a conformation, in which the threonine hydroxyl group can transfer its proton to the histidine base and in which it is positioned for nucleophilic attack on the adjacent peptide carbonyl group at the same time (59 frames for L1 vs. only 1 for B2, Table 2). In the L1 simulations with HSE (His N_ε_ is protonated, Table 2) suitable hydrogen bonding geometries between H837 and T839 are much more frequently observed in comparison to the HSD simulations (Table S4). The quite frequent occurrence of the H837-T839 hydrogen bond in the L1 receptor is due to the high occupancy of the histidine p80 rotamer (χ_1_ = +60°) in the simulation (Table 2). In this rotamer the N_δ_ of the histidine base is positioned for hydrogen bonding interaction to the Ser/Thr nucleophile when this adopts m or p rotamers. Whereas the m rotamer shows even higher occupancy in the B2 simulation (Fig. 5, Table 2), the low occupancy of the p80 rotamer of H912 causes the rare occurrence of the H910-S912 H-bonding contact in B2 and might contribute to the cleavage deficiency of this receptor. We suspect that the different environment of the imidazole side chain of the histidine base in L1 and other cleavage-competent receptors is responsible for ensuring a high occupancy of the p80 rotamer. As noted earlier, the most prominent differences are the T-shaped π-stacking interaction with a phenylalanine or tyrosine residue in place of S876 and the N893G mutation (Fig. 4B).

To test the influence of the conserved glycine and phenylalanine residues on the conformation of the GPS, we analyzed MD simulations of a S876G/N893G variant of the B2 HormR-GAIN domains (Table 2). Indeed, in two of the three replicas, favorable geometries for nucleophilic attack of S912 are observed much more often than for the wild-type receptor (59 frames for L1 compared to 1 frame for B2). Interestingly, while the frequency of favorable θ angles is lower in the L1 simulations, hydrogen bonding interactions between H910 and S912 occur much more often (10627 frames in the L1 simulation compared to 63 frames in the B2 simulation). Concerning both criteria, the S876G/N893G variant is located between wild-type B2 and L1 GAIN domains.

We inspected the MD frames of L1 simulations that have favorable positions of the serine nucleophile and histidine base (Table 2) manually and noted that in some of these structures a distorted ω angle was marked for the scissile peptide bond, in contrast to all residues in the GPS environment. A systematic analysis of the ω torsion angle C_α_-C-N-C_α_, which describes the planarity of the peptide bond, showed that strained conformations of the scissile bond (quantified by having more than 25° deviation from planarity) were observed in 17 (28.8%) of the 59 “GPS-active” conformations listed in column 4 of Table 2 for the L1 GAIN domain. This compares to 1.2 % strained conformations of this peptide bond observed in all structures of the L1 MD simulations. Thus, strained conformations are strongly enriched when the His and Ser residues of the GPS are in a reactive position. A plot of the distortion of the ω angle against the deviation of the position of the Ser nucleophile from the optimal 90° angle for attack on the peptide carbonyl group shows that favorable orientations indeed correlate with strained peptide bonds (Fig. S17). This strain of the ground state structure favors nucleophilic attack on the peptide carbonyl group. The scissile peptide bond is however not unique in comparison to other residues of the GAIN domain concerning the deviations of the ω angle from planarity (Fig. S17C). As noted before, the main chain conformation of the Ser/Thr residues is also strained concerning the φ,ψ-angles observed in the B2 and B3^4^ GAIN domain crystal structures. In the energy minimized AlphaFold model of L1, the nucleophilic T838 is also an outlier in the Ramachandran plot. It appears likely that the strained main-chain conformation at T838 contributes to the formation of strain on the scissile peptide bond to favor autoproteolysis.

Taken together, the typical cleavage-competent GAIN conformation is characterized by deprotonation of the S/T hydroxyl nucleophile by the N_δ_-atom of the GPS histidine in the p80 rotamer (Fig. 6A). Nucleophilic addition to the neighboring peptide carbonyl group is supported by strain on the peptide bond. The p80 rotamer is stabilized by a T-shaped π-π interaction with a phenyl side chain (F831 in L1) and a hydrogen bond to the carbonyl oxygen of the peptide bond to the highly conserved glycine-cysteine residues N-terminal to cysteine bridge 2 (CC2, Fig. 6A). Further hydrogen bonds of the main-chain of this region upstream of the GPS additionally stabilize the GPS-active state in this model. These interactions were observed in all MD conformations of the L1 simulation, that are considered active concerning the first step of the autoproteolysis reaction.

**Figure 6:**
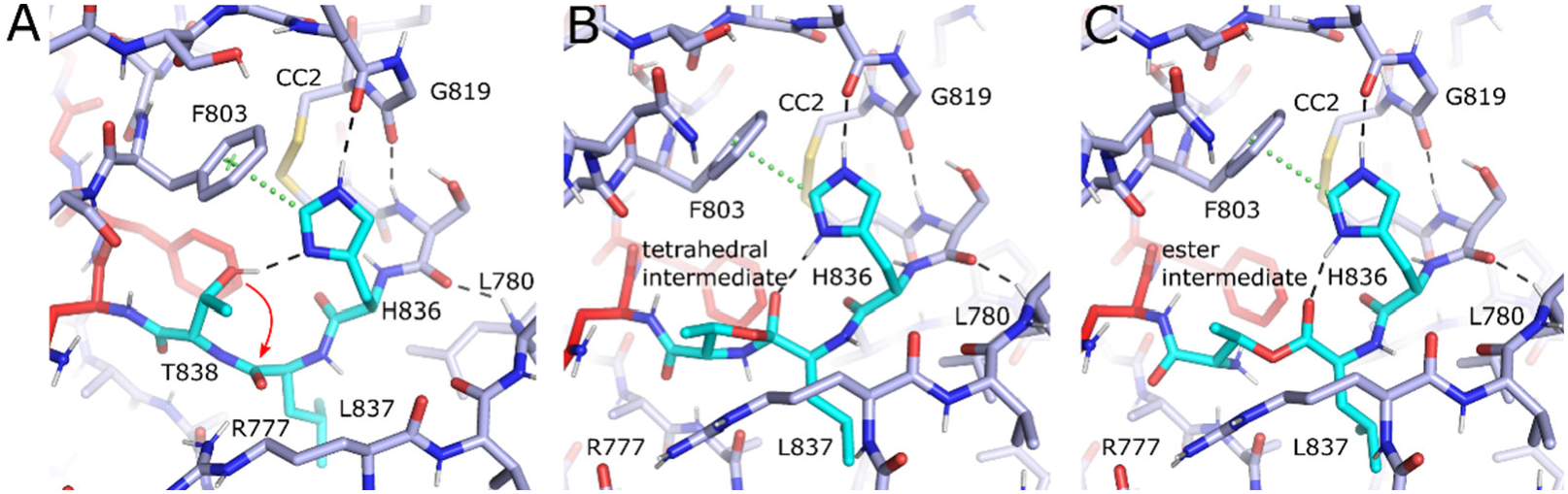
Models of the catalytic mechanism of GPS hydrolysis. (A) Model of the GPS in a favorable geometry for deprotonation of T838 by the H836 base and subsequent nucleophilic attack of the hydroxylate on the carbonyl carbon. The model was obtained from a representative frame of the MD simulation of the L1 GAIN domain. H836 adopts the p80 rotamer (χ_1_ = 61°) and T838 the p rotamer (χ_1_ = 59°). (B) Model for the tetrahedral intermediate or transition state **2** (see Figure 1). (C) Model of the ester intermediate. R777 is positioned in the models constructed from this MD frame to participate in the catalytic mechanism.

To analyze interactions that might catalyze further steps of the autoproteolytic mechanism, we generated a model of the tetrahedral intermediate or transition state formed after nucleophilic attack of the Ser hydroxyl (Fig. 6B). In this model the protonated catalytic histidine might stabilize the negatively charged intermediate by hydrogen bonding interactions to the oxygen atom that results from the carbonyl oxygen of the peptide bond. Although this model is the result of only energy minimization and the conformational flexibility of this intermediate has not been investigated here by MD simulations or other *in silico* techniques, it appears unlikely that the main chain can rearrange such that the histidine would be positioned to directly protonate the leaving nitrogen atom in the next catalytic step, as suggested in the catalytic model shown in Figure 1. Since also no other protein residue of the conserved core residues of the GAIN domain (Fig. 4B and 6A) is positioned for this protonation step, we assume that a solvent water molecule provides a proton upon expulsion of the nitrogen atom. It is interesting to note that the side chain of R777, which is part of the flexible flap 1, is in the direct vicinity of the scissile peptide bond in some frames of the L1 simulation. The R777 side chain is oriented away from the GPS into the solvent in the crystal structure of the L1 GAIN domain^4^.

After the nitrogen atom has left, the ester intermediate is formed (Fig. 6C). Besides the histidine base there is no obvious residue in the immediate environment of the GPS that could support hydrolysis of the ester intermediate by stabilization of the transition state (other than R777 of the flexible flap1). The basic primary amine resulting from the released peptide nitrogen likely acts as a general base to deprotonate a water nucleophile for attack on the ester carbonyl as the last step of the autoproteolytic mechanism. Deeper mechanistic understanding of these steps requires an analysis of the energy landscape and flexibility of the two intermediates.

### The full-length B2 receptor is not cleaved at the GPS upon expression in HEK293T cells

Even though it was shown for L1, G1, G3, and G6 that the GAIN domain is sufficient for self-cleavage^4–7^, a possible cause of the lack of GPS cleavage for the ectodomain constructs of B2 may be the lack of cleavage-modulating factors. For example, we surmised that the presence of the 7TM domain may aid in activation of the GAIN domain for the autoproteolysis reaction. We tested cleavage in HEK293T cells with different constructs and detected only bands of the uncleaved receptor with apparent molecular masses that correspond to the monomeric and dimeric receptor (Fig. 7A). A band observed at ~95 kDa may correspond to an N-terminal fragment that results from additional proteolytic cleavage, akin to the release of Vstat-40 from *h*B1^32^, e.g. at the long disordered loop 770-810. The fact that this fragment is also present for the H910A and S912A variants rules out that it is derived from GPS cleavage.

**Figure 7:**
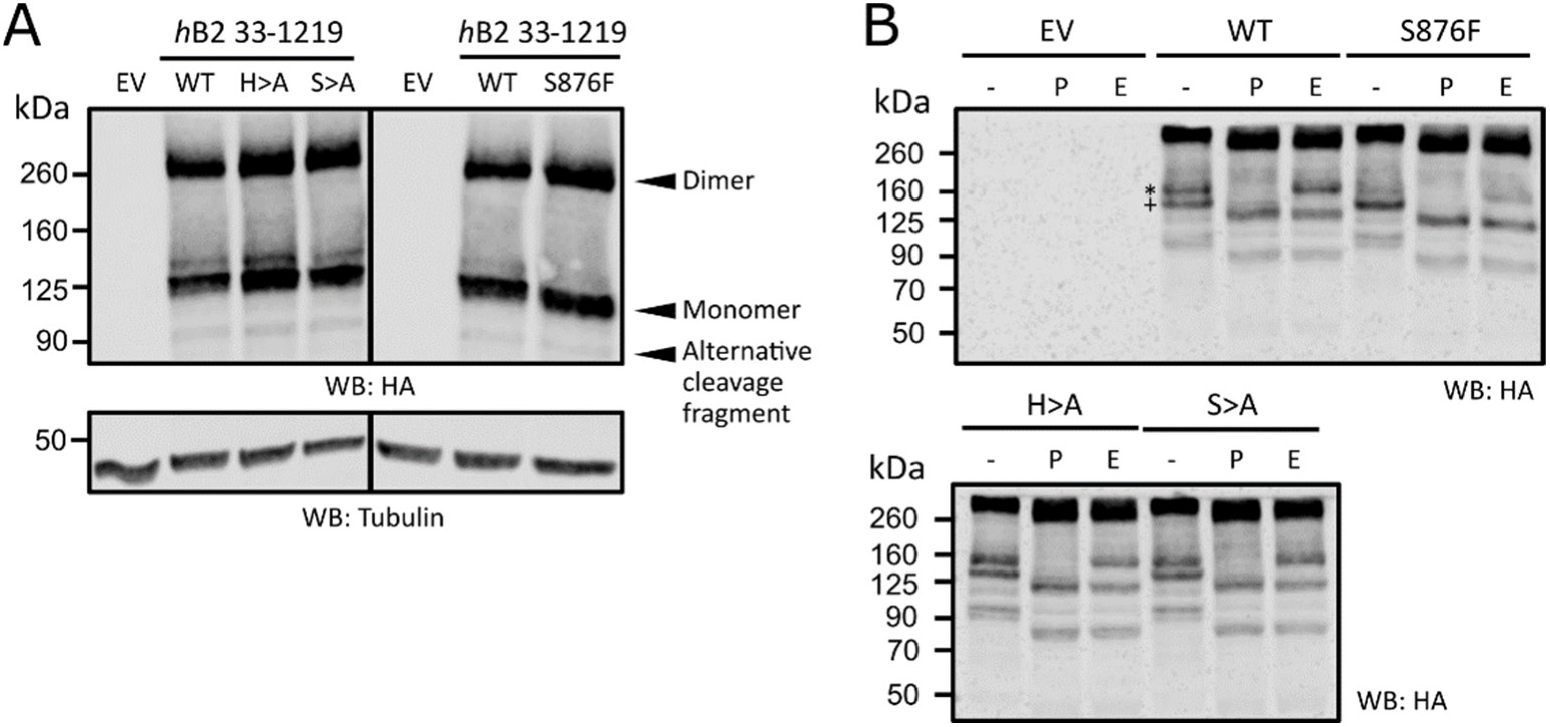
Autoproteolysis is not observed for the full-length *h*B2. HEK293T cells were transiently transfected with indicated mutated constructs of *h*B2 residues 33-1219. (A) Western blotting of an SDS-PAGE showed the expressions of *h*B2 and respective mutants targeted against the N-terminal HA tag of the receptor. 30 μl of lysate samples were used for analyses. Tubulin served as loading control. “H>A” denotes the H910A mutation, “S>A” the S912A mutation and “EV” the empty vector control. (B) Western blotting of SDS-PAGE of *h*B2 lysates treated with PNGase F (P), Endo H (E), or without any enzyme treatment (-). Endo H-resistant and sensitive bands of monomeric *h*B2 indicated by “*” and “+”, respectively. Calculated molecular masses and further details see Figure S19.

### Evolutionary conservation of the B2 GAIN domain

We analyzed the conservation of the amino acid sequence of the human B2 receptor by a comparison to 148 mammalian orthologs to indicate functional roles of specific protein residues for folding, stability and in particular for GPS hydrolysis (Fig. 8). Flaps 1 and 2 and also the long disordered loop of residues 770-810 (Fig. S20) show little sequence conservation. We did not find any residues in the environment of the GPS that are highly conserved to suggest a catalytic function. Notably, also the histidine base position (H910) shows significant variance. In detail, the −2 position of the GPS sequence contains His (41.1%), Arg (36.4%), Gln (11.9%), Lys (9.3%), and Tyr (1.3%) residues. The +1 residue contains a serine in 95.4% and an alanine in 4.6% of the sequences. Thus, in the majority of the B2 orthologs the GPS sequence indicates cleavage resistance. In L1, the GPS and also the neighboring flap 1 region is much higher conserved compared to B2 and B3 (Fig. 8). However, the R^-2^ residue at the GPS of B3 is also strictly conserved with one exception for the fish *Tetraodon nigroviridis*, although this residue has no obvious function for GPS cleavage or *Stachel* dissociation. It may have a structural role in the formation of a salt bridge with a strictly conserved aspartate residue (D823 in human B3).

**Figure 8:**
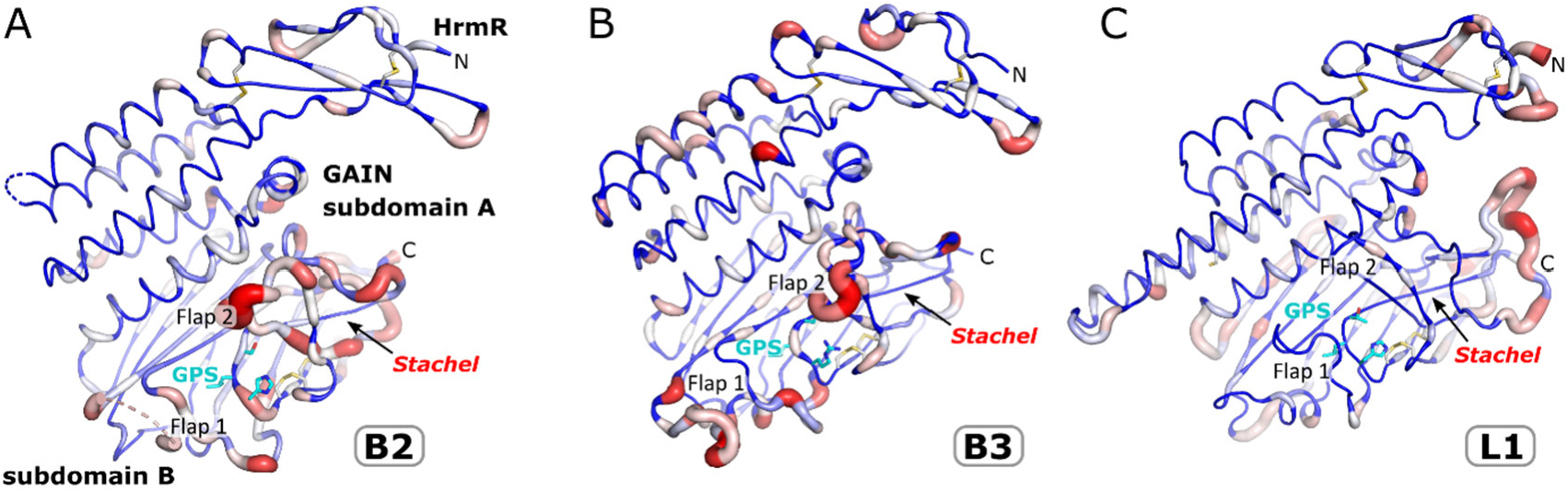
Conservation of the B2, B3, and L1 HormR and GAIN domains. Sequences of the HormR and GAIN domains of 148 (B2), 137 (B3), and 167 (L1) mammalian species were aligned and the Shannon entropy was calculated for each residue, which quantifies the variability of the sequence positions. Residues of high conservation are shown as blue thin tube regions and residues of high sequence variation as red thick tube regions.

### Cleavage competence of ADGRB3 can be restored by a double mutation

Whereas trials to restore cleavage competence of the *h*B2-HG construct by a S876F/N893G double mutation failed, we were able to activate ADGRB3 for self-cleavage (Fig. 9). This receptor features an RLS motif at the GPS (with R866 at the base position) and the conserved aromatic residue that forms a T-shaped π-π interaction with the histidine base in cleavage-competent receptors (Fig. 5B) is L821 in B3. An R866H and L821F or L821Y double mutation restores the general base and its edge-π interaction for stabilization of the p80 rotamer and the mutant receptor displays weak but significant cleavage activity (Fig. 9). The R866H mutation alone results in lower auto-proteolytic activity. Vice versa, the naturally cleavage-competent L1 receptor shows reduction in its autoproteolytic activity if F803 (the residue for the edge-π interaction with the histidine base H836) is replaced by a leucine or alanine, whereas full cleavage competence is retained for F803W and F803Y mutations. These findings demonstrate the importance of the edge-π interaction of the histidine base for self-cleavage activity. The residual cleavage competence of the F803A and F803L variants of L1 as well as the limited reactivation of B3 in the R866H/L821F variant shows that additional factors further contribute to cleavage competence.

**Figure 9:**
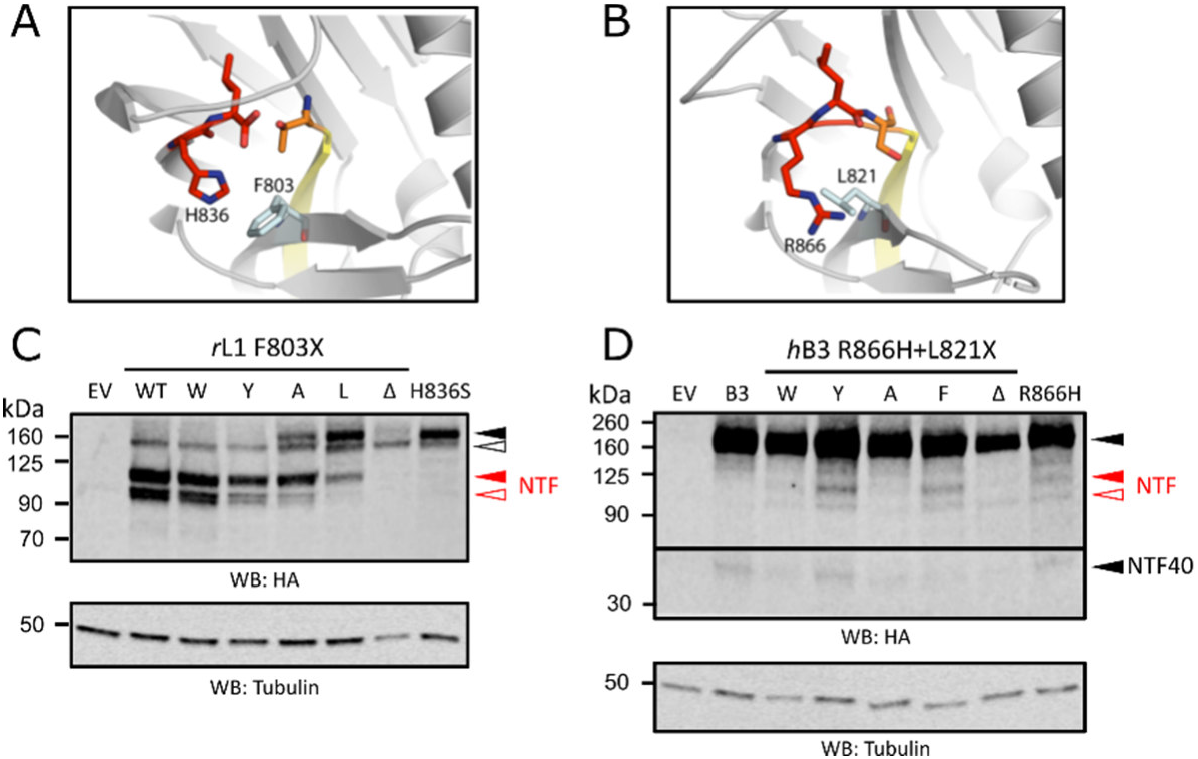
Edge-π interaction between the His^-2^ base and surrounding residues is important for GPS cleavage of some receptors. (A) Edge-π interaction between H826 and F803 observed in the crystal structure of rat L1 (pdb id 4dlq)^4^, which is a naturally GPS-cleavable receptor. (B) These residues correspond to R866 and L821, respectively, in the cleavage-resistant *h*B3 (crystal structure 4dlo^4^). (C,D) HEK293T cells were transiently transfected with constructs containing mutations causing either a disruption of edge-π interaction in rL1 (C), or a reintroduction of edge-π interaction in *h*B3 (D). GPS cleavages of the mutants were analyzed by WB using 30 μl of cell lysates, targeted against the N-terminal HA tag of the receptors. Tubulin served as a loading control. Bands representing uncleaved receptor and NTF are indicated in black and red triangles, respectively. Fully (N- and O-glycosylation) and immaturely (no O-glycosylation) glycosylated populations of rat L1 are indicated in closed and open triangles, respectively. “NTF40” denotes an N-terminal fragment of ~43 kDa apparent molecular mass indicating additional processing of *h*B3, similar to *h*B1^32^ and *h*B2 (Fig. 7A). For further details including calculated molecular masses, deglycosylation experiments and interpretations see Figure S21.

### Dimer structure of the *h*B2-HG construct

As described before, BN-PAGE and DLS analysis indicated dimer formation for all three expressed constructs. In the crystal structure of *h*B2-HG, we identified four different candidates for a dimeric arrangement via the interfaces between symmetry mates within the crystal lattice (Table S5). Two of the identified dimers (#1 and #3) possess a closed C2-symmetry (Fig. S22), whereas the other dimers are asymmetric. These two dimers had similar predicted free energies of interaction. We therefore collected small angle X-ray scattering (SAXS) data to distinguish between the two possible dimer structures.

SAXS data was recorded for a dilution series of *h*B2-HG between 0.5 mg/mL and 10 mg/mL (Fig. S23). In the investigated concentration range the radius of gyration increases and the maximum of the Kratky plot is shifted away from a value close to the theoretical value of a solid sphere to larger values, in agreement with the elongated dimer structures compared to the more compact monomer fold. For a comparison of the theoretical and experimental scattering curves, the crystallographic models for the monomer and the two symmetric dimers were complemented by high mannose N-glycans and by the loops and terminal amino acids, that were not resolved in the crystal structures. Ensembles containing either only monomer or dimer models or a mixture of monomer and dimer models were fitted to the SAXS curves (Fig. S23, Table 3). The monomer fraction was reduced from 85% at 0.5 mg/ml to 66 % at 10 mg/ml. The limited concentration range that could be achieved by SAXS measurements does not allow for an accurate determination of the K_D_-value, but a rough estimate of K_D_ ≈ 400 μM was obtained. This rather transient interaction is in agreement with the broad bands observed in BN-PAGE experiments. We assume that larger ECR constructs form dimers with higher affinity, as indicated by the sharper BN-PAGE bands. TSP domains are involved in homodimer formation^33^. The χ^2^ values at higher protein concentrations show that dimer #3 is present in solution (Table 3, Fig. S22). This dimer is formed by interactions between subdomain A of the GAIN domain. In both dimer models the HormR domain is located at the same site as the end of the *Stachel* sequence, such that the HormR domains are placed close to the cell membrane (Fig. S24).

**Table 3.** Radius of gyration and goodness-of-fit (χ^2^) parameters derived from the SAXS data analysis. The best fits are indicated by χ^2^-values close to 1. The numbers in brackets indicate the molar fraction of the monomer in the ensemble.

| c [mg/mL] | R<sub>g</sub> [nm]<br>(Guinier) | $\chi^2$ (monomer molar fraction) | | | | |
| --- | --- | --- | --- | --- | --- | --- |
|  |  | Monomer | Dimer #1 | Dimer #3 | Monomer/<br>Dimer #1 | Monomer/<br>Dimer #3 |
| 0.5 | 3.08 | 3.58 | 4.06 | 4.59 | 0.98 (0.839) | 0.78 (0.848) |
| 1.0 | 3.08 | 9.55 | 15.95 | 16.24 | 1.65 (0.859) | 0.85 (0.858) |
| 2.5 | 3.19 | 53.64 | 55.56 | 40.27 | 7.94 (0.830) | 0.91 (0.826) |
| 5.0 | 3.31 | 148.41 | 126.12 | 45.36 | 27.11 (0.796) | 1.59 (0.759) |
| 10.0 | 3.47 | 107.09 | 57.79 | 11.83 | 14.47 (0.766) | 1.64 (0.659) |

## Discussion

Expression of the ectodomain of the human ADGRB2 receptor showed that it forms a dimer and is resistant to autoproteolysis at the GPS, despite the presence of a canonical HLS motif. We could crystallize the HormR-GAIN domains of ADGRB2 and determined the crystal structure of this aGPCR GAIN domain in the uncleaved state. After structure determination of the HormR-GAIN domains of ADGRB3^4^, this is the second structure of an uncleaved aGPCR GAIN domain, whereas four structures are available for GPS-cleaved GAIN domains (L1, G1, G3, G6)^4–7^. A comparison of the B2 and B3 GAIN domains with AlphaFold models of cleavage-competent aGPCR indicated that the fold of the GPS sequence is rather similar and that the immediate environment of the GPS in the GPS-active receptors does not contain any conserved residues besides the GPS sequence itself, that are capable of acid-base catalysis, which would be a typical situation for a proteolytic enzyme. On the contrary, the GPS environment lacks such residues and it displays a surprisingly diverse environment in particular in the form of two flaps, that are flexible and flank the GPS groove, as noted in previous work^16^.

However, there are also structural features that are conserved in cleavage-competent GAIN domains. They contain a highly conserved phenylalanine that interacts with the histidine side chain of the HXS/T GPS sequence via T-shaped π-stacking. Together with a hydrogen bond to the main chain carbonyl group of the residue preceding the highly conserved Gly-Cys motif in GPS-active aGPCR, this interaction positions the histidine side chain to accept a proton from the serine or threonine nucleophile. The cysteine of the Gly-Cys motif forms a disulfide bridge that is present in most aGPCR GAIN domains. MD simulations of the cleavage-competent L1 GAIN domain in comparison to the GPS-inactive B2 GAIN domain demonstrated the importance of these interactions for the formation of cleavage-competent conformations, in which the histidine base, the serine/threonine nucleophile and the electrophilic carbonyl carbon are in a favorable position for the first step of the catalytic mechanism (Fig. 6A). The formation of this conformation further results in the formation of strain on the scissile peptide bond, which likely favors nucleophilic addition of the alcohol to the peptide carbonyl group.

Mutation of the π-stacking phenylalanine to an alanine or leucine strongly reduced the autoproteolytic activity of L1 whereas the cleavage-incompetent B3 GAIN domain could be activated by a double mutation that installed the phenylalanine-histidine π-stacking pair at the GPS (Fig. 9). The residual activity of the L1 mutant, the only partial activation of B3 and the failure to activate the B2 GAIN domain via an S876F/N893G mutation indicates that other factors may contribute to maximal autoproteolytic activity. We also generated models of the tetrahedral intermediate and the ester intermediate of the catalytic mechanism (Fig. 1) and found no interaction partners that would be positioned to function as an oxyanion hole as known from serine proteases. The water nucleophile for release of the ester intermediate is likely activated via deprotonation by the basic primary amine group released from the peptide bond. These models show that the histidine cannot protonate the nitrogen leaving group of the tetrahedral intermediate directly. Further information on these steps of the catalytic mechanism requires a more detailed analysis of the conformational landscape and energy. The differences between the GPS with its environment and serine proteases also reflect the vastly different reactivity. Whereas serine proteases can have turnover numbers of around 100 s^-1^, the autoproteolytic reaction needs to be catalyzed for only one turnover on the timescale of minutes to hours.

*Cis*-autoproteolysis, where a serine, threonine or cysteine nucleophile attacks the penultimate peptide bond followed by N→(O,S) acyl shift and formation of the ester intermediate, has also been observed in other proteins ^34^, including N-terminal nucleophile (NtN) hydrolases^35,36^, the pantetheine hydrolase ThnT^37^, Nup98^38^ or the SEA domain^39,40^. In mechanistically related non-hydrolytic proteins, the (thio)ester intermediate is attacked by nucleophiles other than water, such as in inteins^41^, hedgehog proteins^42^ or pyruvoyl enzymes^43^. Despite the common catalytic features, these proteins employ significantly different additional strategies to catalyze the self-cleavage reaction (Table S6). A common structural feature is that the scissile peptide bonds are solvent exposed and positioned in loop regions next to a β-strand. However, the conformation and environment of the processed peptide region is diverse. A factor, that appears to be more prominent in *cis*-autocleavage compared to serine proteases or other enzymes, is the use of strain to destabilize the ground state. Distortions from ideal geometry have been observed for the scissile peptide bond and for adjacent torsion angles for many independent systems (Table S6). Strain or conformational energy of adjacent bonds is likely temporarily transferred to the scissile peptide bond by protein dynamics. Distortion of the peptide bond results in a drastic increase of the pK_a_ of the nitrogen such that it may be protonated even before attack of the nucleophile^44^. The role of strain in *cis*-autoproteolysis has been especially well studied for SEA domains, resulting in a model for the catalytic mechanism where substantial energy of protein folding is invested to generate a strained flexible pre-cleavage ground state structure, which is poised for nucleophilic attack in the absence of a catalytic base and oxo-anion hole^39,45^. Remarkably, even removal of the hydroxyl nucleophile by mutation to an alanine resulted in residual site-specific cleavage activity.

Analysis of the GPS sequences of B2 homologs demonstrated that the majority of these have no cleavage-competent GPS sequence. In comparison, the GPS of L1 is strictly conserved as a H-L-T sequence motif while B3 homologs contain a highly conserved R-V/L-S motif. The human B2 receptor has a canonical HLS sequence at the GPS, but the environment of the GPS, in particular the lack of a phenylalanine at position 876 and of a glycine at position 893 distinguishes this receptor from autoproteolytically active GAIN domains. As the S876F/N893G double did not reinstate cleavage activity, further yet unknown structural features inactivate the B2 receptor for GPS cleavage. GPS autoproteolysis was also not observed for the full-length protein expressed in HEK293T cells (Fig. 7). Taken together, these findings suggest that the B2 aGPCR signals without GPS cleavage, as it has also been shown for the naturally autoproteolysis-resistant receptors ADGRG5/GPR114^24^ and ADGRC3/CelsR3^46^ This finding is in agreement with previous studies demonstrating no GPS cleavage for B1, B2 and B3 for expression in HEK293T cells. In contrast, GPS-cleavage of B2 was concluded in a previous study^47^. In agreement with our results, the majority of the B2 receptor obtained from recombinant expression in U87-MG cells was not cleaved, but the observation of several fragments in a western blot of this cell line and of mouse hippocampal tissue was interpreted as CTF fragments in this study. Likewise, fragments corresponding in size to a CTF fragment were observed in mouse brain lysates and brain immunoprecipitates with anti-BAI1 and anti-BAI3 antibodies raised against cytosolic protein regions^4^. These fragments obtained from endogenously expressed receptors might correspond to the CTFs generated by GPS cleavage. Alternatively, they might be expressed from alternative promotor regions located before the coding region of the CTF. For B3, as well as for 67% of all aGPCR genes, mRNA transcripts of CTFs without or with a short N-terminal extension were characterized^48^. Therefore, the origin of putative CTF fragments of ADGRB2 and ADGRB3 detected in native tissue requires more detailed characterization.

GPS cleavage influences the most important function of the GAIN domain, the release of the *Stachel* sequence for binding to the 7TM domain. As the Ser/Thr residue of the GPS is the first residue of the *Stachel*, the GPS environment (Fig. 6A, 4B) has been optimized by evolution not only to enable or disable GPS cleavage but also to enable exposition of this sequence^49^, likely in response to mechanical stimuli for many aGPCR^50–58^. As noted since characterization of the first GAIN domain structure^4^, the apparently tight association of this sequence as a central β-strand in the GAIN subdomain B (Fig. 10A) is puzzling, also considering the high thermal stability of GAIN domains. The B2 HormR-GAIN domain has a melting point of 76.6±0.08°C (Fig. S6) and similar values have been observed by us for other GAIN domains. However, a similar rearrangement of a central β-strand is also observed in other proteins, such as the serpins. In these proteins, proteolytic cleavage of a loop region results in an insertion of part of this loop into a β-sheet^59^. We speculate that the observed flexibility of the flanking loop regions observed in MD simulations in this and previous^16^ studies contributes to the dynamic process of *Stachel* release. AlphaFold predictions^60^ of all aGPCR indicate no significant interaction between the GAIN domain and the 7TM domain that would result in a stable association of these two domains. Furthermore, the GAIN domain could so far not be visualized in cryo-EM studies of GAIN-7TM constructs^17,19,18,20,22,21,23^, likely due to flexible linkage of the two domains. This situation favors a “one-and-done” signaling mechanism (dissociation model), in which *Stachel* binding to the 7TM domain is preceded by dissociation of the GAIN domain, NTF release^49^ and permanent receptor activation, only terminated by internalization^61^. In contrast, GPCR signaling in cleavage-deficient aGPCR may operate also via a reversible signaling mode, in which the *Stachel* sequence exists in an equilibrium between 7TM- and GAIN-bound states (non-dissociation model). This process likely requires the unfolding of the four terminal β-strands of GAIN subdomain B, whereas the residual part of this domain might maintain a folded state (Fig. 10C), supported by the high thermal stability of this part of the GAIN fold. The structural flexibility of the loops flanking the GPS might support this unfolding step^16^. It is interesting to note, that the highly conserved disulfide bridges (CC1 and CC2, Fig. 10A) link all four terminal β-strands and might ensure fast refolding kinetics in addition to the stabilization of the GPS environment for autoproteolysis in cleavage-competent receptors. An alternative scenario could be that the *Stachel* sequence is prebound in the 7TM domain binding pocket and isomerizes after mechanoactivation or ligand binding^52^ as found in glycoprotein hormone receptors^62^.

**Figure 10:**
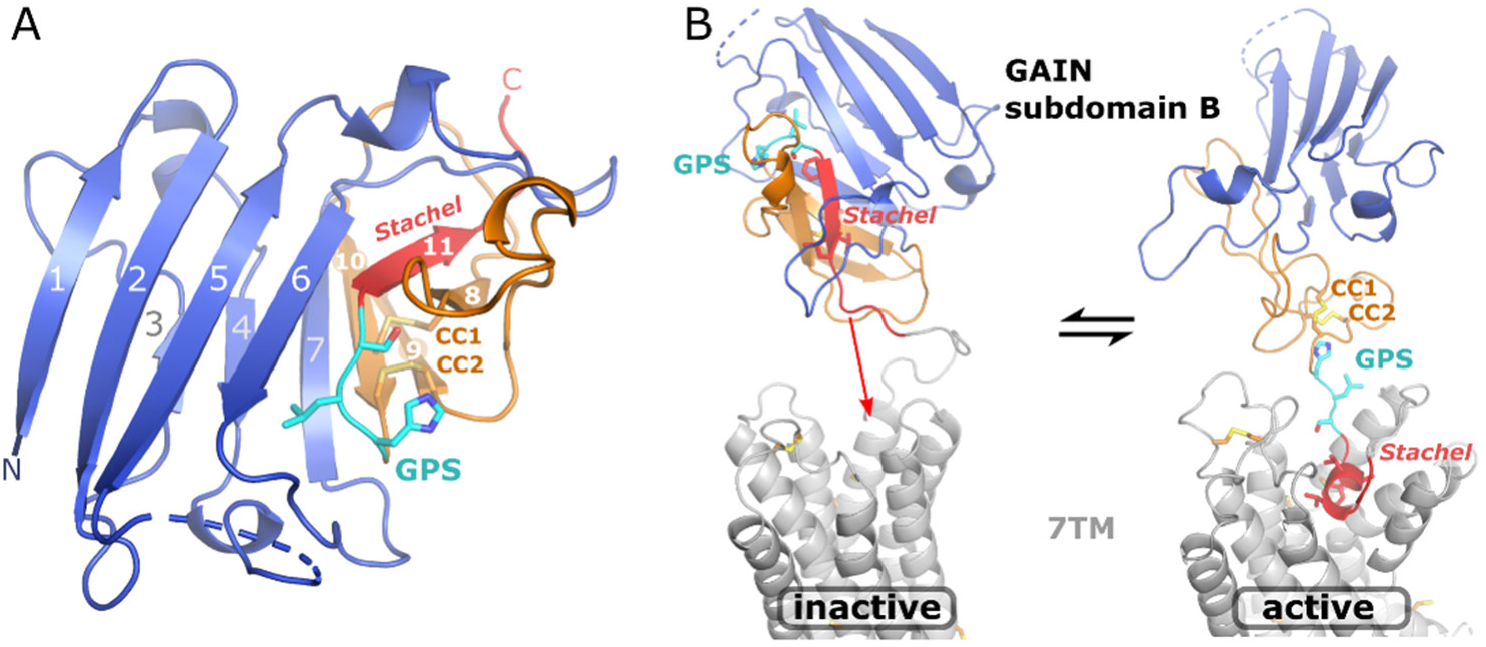
Fold of GAIN subdomain B and a model of *Stachel* dissociation. (A) Fold of subdomain B of the *h*B2. The region comprising the terminal four β-strands 8-11 would probably unfold as a result of dissociation of the *Stachel* sequence from the GAIN domain. Two highly conserved disulfide bridges (CC1 and CC2) link β-strands 8-10 and they are well conserved in most but not all aGPCR. β-strands 7-11 are the conserved region denoted as GPS before characterization of the GAIN domain structure^4^ (B) Illustrative hypothetical model of an equilibrium of *Stachel* association with the GAIN and 7 TM domains. The orientation of GAIN and 7TM domains is arbitrary. This figure is shown to illustrate the length of the linker between the β-strand of the *Stachel* (when bound to the GAIN domain) and the 7TM domain and the conserved mode of binding of the *Stachel* to the 7TM domain as observed in cryoEM structures. CC1 and CC2 might help to ensure fast refolding kinetics of the terminal four β-strands of the GAIN domain.

Our understanding of GPS cleavage or non-cleavage as well as *Stachel* release from the GAIN domain needs to be extended by studies probing the influence of further factors contributing to these highly dynamic processes. In particular, the reversibility of *Stachel* dissociation and its influence on GPCR signaling is a fascinating aspect of future research. For GPS-cleaved receptors, it remains to be shown, if yet unknown interactions with the 7TM domain or the stability of interactions in the network of the extracellular matrix maintain the GAIN domain in the vicinity of the 7TM domain for re-association with the *Stachel* agonist.

## Materials and Methods

### Cloning

DNA encoding the ECR of human ADGRB2/BAI2 (aa 1-921) was synthesized by Thermo Fisher GeneArt. The desired inserts (*h*B2-ECR residues 33-921, *h*B2-C 33-295, *h*B2-THG 296-921, *h*B2-HG 528-921) were subcloned into a pHLsec vector^63^ for transient transfection and small-scale expression tests, and subsequently into a modified PiggyBac transposon vector^64^ for stable transfection and large-scale expression (Fig. S25). For all cloning steps a classical approach using PCR and restriction endonucleases was used (Table S7). The final expression constructs included a C-terminal enteropeptidase recognition site followed by a Twin-Strep-tag^®^ (TST). *h*B2-ECR and *h*B2-HG included an additional N-terminal hemagglutinin antibody-tag (HA-tag).

### Small-scale expression tests und generation of stable cell lines

HEK293S GnTI^-^ cells were cultured in Dulbecco’s Modified Eagle Medium (DMEM; ThermoFisher Scientific, cat# 21969035) supplemented with 10 % (v/v) fetal bovine serum (FBS; Gibco cat# 10437028, Life technologies), 1 % (v/v) GlutaMAX (cat# 35050038, Life technologies) and 1 % (v/v) MEM non-essential amino acids (cat# 11140035, Life technologies) and kept in a humidified incubator at 37°C balanced with 5 % CO_2_. DMEM containing only 2 % (v/v) FBS was used during transient protein expression and serum-free DMEM was used to prepare transfection mixtures. Our DMEM formulation for storing cells in liquid nitrogen contained 20 % (v/v) FBS and 10 % (v/v) DMSO (cat# D8418, Sigma Aldrich). All reagents were warmed up to 37°C prior to usage.

HEK293S GnTI^-^ cells were grown to approximately 90 % confluency on 24-well tissue culture plates (cat# 11391704, Fisher Scientific) before the medium was exchanged for serum reduced DMEM and the transfection mixture was added. The transfection mixture contained ~500 ng of the plasmid carrying the target sequence. Six different transfection conditions (Table S8) were tested by adding 0, 100, or 500 ng of the pAdVantage vector, which encodes adenoviral virus associated I RNA, and 750, 900, or 1500 ng polyethylenimine (cat# 408727, Sigma Aldrich). After incubation for 15 min at room temperature the mixture was added to the cells. The addition of 4 μM valproic acid 3 h after initial addition of the transfection mixture was tested for its effect on the expression yield. The medium was harvested and analyzed via western blot after 5 days.

The PiggyBac Single Promotor vector (cat# PB531A-1, System Biosciences) was modified in house to replace the gene encoding the red fluorescent protein (RFP) by a gene for green fluorescent protein (GFP) expression. In addition, the multiple cloning site was modified for subcloning via the *Eco*RI and *XhoI* restriction sites from the pHLsec vectors. Stable cell lines were generated by cotransfection with the Super PiggyBac Transposase Expression Vector (cat# PB210PA-1, System Biosciences) followed by selection via addition of 5 μg/mL puromycin (cat# ant-pr-1, InvivoGen) over 10 days. Cells were then allowed to recuperate for approximately seven days before they were transferred into cryo-storage until further use.

### Large-scale protein expression and purification

Stably transfected HEK293S GnTI^-^ cells were retrieved from cryo-storage and gradually scaled up over three passages until they were seeded in six roller bottles (cat# 681670, Greiner Bio-One), where they were maintained for 10 days with an additional medium exchange after 5 days. The conditioned medium from all passaging steps and the roller bottles was collected and pooled for purification.

1 mL BioLock biotin blocking solution (cat# 2-0205-050, IBA Lifesciences) was added per 1.5 L of conditioned medium and incubated for 2 h gently stirring at 4°C. The medium was then centrifuged for 30 min at 18,600 g and the supernatant filtered through a 0.22 μm polyethersulfone membrane (cat# 514-0332P, Avantor). Prior to affinity chromatography, the medium was concentrated approximately 15-fold, using a hollow fiber cartridge with a surface area of 4,800 cm^2^ and a nominal molecular weight cutoff (NMWC) of 10 kDa (GE Healthcare). All chromatography procedures were carried out at 4°C using either an ÄKTA express or ÄKTA pure chromatography system (GE Healthcare). 1 mL StrepTrap^TM^ HP (cat# 28907546, Cytiva) was used for affinity chromatography and a HiLoad^TM^ 16/600 Superdex^TM^ 200 pg (cat# GE28-9893-35, Sigma Aldrich) for size exclusion chromatography (SEC). Before injection into the SEC column, samples were concentrated to a volume of approximately 2 mL using Amicon^®^ Ultra centrifugal filters with a NMWC of 10 kDa (cat# UFC801024, Merck Millipore).

### Mass spectrometric analysis of the furin cleavage fragments

Two gel plugs were cut from each Coomassie stained band with the help of an ExQuest™ spot cutter (Bio-Rad Laboratories GmbH, Feldkirchen) and placed in a 0.5 mL Eppendorf reaction tube. For destaining, gel plugs were washed three times with 100 μL of a 30 % (v/v) acetonitrile (cat # 1204102BS, Bisolve B. V., Valkenswaard, Netherlands) in 50 mmol /L ammoniumbicarbonate, dehydrated with 100 μL acetonitrile and rehydrated with 25 μL trypsin solution, containing 5 ng/μL trypsin (cat # 37283, SERVA Electrophoresis GmbH, Heidelberg) in 3 mmol/L ammoniumbicarbonate (cat # 09830, Merck KGaA, Darmstadt). After 4 h incubation at 37 °C, peptides were extracted by incubating the gel plugs with 20 μL 60 % (v/v) acetonitrile with 0.1 % (v/v) formic acid (cat # 6914143BS, Bisolve B. V., Valkenswaard, Netherlands) and 20 μL acetonitrile. The combined extracts were dried in vacuum centrifuge and peptides were re-dissolved in 20 μL 3 % (v/v) acetonitrile and 0.1 % (v/v) formic acid.

Analysis was performed on a nanoAcquity UPLC system (Waters GmbH, Eschborn) coupled on-line to a LTQ Orbitrap XL mass spectrometer with a Nanospray Flex Series ion source (Thermo Fisher Scientific, Bremen). 1 μL peptide solution was injected on a Symmetry C18 column (particle size 5 μm, 180 μm internal diameter (ID), 2 cm length). Peptides were trapped with a flow rate of 5 μL/min 3 % eluent B (acetonitrile containing 0.1 % (v/v) formic acid) where eluent A is 0.1 % (v/v) aqueous formic acid for 6 minutes followed by separation on a BEH C18 column (1.7 μm particle size, 75 μm ID, 100 mm length) at a flow rate of 0.3 μL/min and a column temperature of 35 °C. Peptides were eluted with linear gradients from 3 to 40 % eluent B in 18.5 minutes and up to 95 % eluent B in 5.5 minutes. Nanospray PicoTip Emitter (New Objective, Littleton) was set at 1.5 kV and transfer capillary temperature to 200 °C. Mass spectra were recorded in *m/z* range from 400 to 2000 in the Orbitrap mass analyzer at a resolution of 60,000 at *m/z* 400. Using DDA tandem mass spectra were acquired in the ion trap with CID mode (isolation width of 2 *m/z* units, normalized collision energy of 35%, activation time of 30 s, default charge state of 2, intensity threshold of 300 counts, dynamic exclusion window of 60 s) for the six most intense ions.

Raw data were processed with PEAKS Studio (version 10.6, Bioinformatics Solutions, Waterloo) using 10 ppm parent mass error tolerance, 0.8 Da fragment mass error tolerance, enzyme trypsin semi specific, 3 miss cleavages, methionine oxidation as variable modification (+15.99 Da) and a custom build database containing the assumed amino acid sequence of the protein construct plus 20,404 other reviewed human protein sequences (loaded 2/7/2023 from https://www.uniprot.org). Results were exported as peptide.pep.xml for library building in Skyline. Quantification relied on the peak areas of the corresponding extracted ion chromatograms (XICs) calculated by Skyline (22.2.0.312 MacCoss, Seattle, WA, USA).

### Autoproteolysis assays of full-length constructs

#### Cloning

For cellular expression analyses of receptors in HEK293T cells, cDNAs encoding *h*B2 33-1219 were first subcloned from pHLsec vector into pcDNA3.1(+) vector before mutagenesis. Mutations on *h*B2^S876F^, *h*B2^H910A^, *h*B2^S912A^, *h*B3^R866H^ and all *r*L1 mutations were made by PCR and sequence-ligation-independent cloning approach (Table S9). *h*B3^R866H/L821X^ was synthesised by Genscript.

#### Cellular expression analyses of receptors

HEK293T cells were cultured in DMEM supplemented with 10% fetal bovine serum and 1% Pen/Strep (Gibco^TM^, cat# 15070063), and maintained in an incubator in 37°C, 5% CO_2_ and saturated humidity. Cells were seeded into 24-well cell culture plate (Greiner, Cat# 662160) one day before transfection, at which point 70-80% confluence was observed. Cells were then transiently transfected with receptor constructs using Lipofectamine^TM^ 2000 transfection system (ThermoFisher, cat# 11668019). Briefly, 400 ng of plasmid in serum-free DMEM was mixed with 1.6μl of Lipofectamine^TM^ 2000 reagent in serum-free DMEM. After 20 minutes, the transfection mixture was added into a well of cells refreshed with full DMEM (10% fetal bovine serum and 1% Pen/Strep). The cells were incubated for one day before lysis with 150 μl triton X-100 buffer (50 mM Tris pH 8.0, 150 mM NaCl, 1% Triton X-100). After removing cell debris, lysates were mixed with Laemmli buffer, followed by SDS-PAGE and WB analyses.

#### Deglycosylation of hB2 33-1219 receptors

HEK293T cells were seeded into 6-well cell culture plates (Greiner, cat# 657160) one day before transfection. Cells were transiently transfected using Lipofectamine^TM^ 2000 transfection system (ThermoFisher, cat# 11668019 similarly as aforementioned, with 1600 ng of plasmid and 6.4 μl of Lipofectamine^TM^ 2000 reagent. Cells were lysed with 500 μl of triton X-100 buffer buffer one day after transfection. For examining N-glycosylation of receptors, 17 μl of the lysates was treated with 500 U PNGase F (NEB, cat# P0704S), 500 U Endo H (NEB, cat# P0702S) or distilled water, supplemented with 2 μl of their respective 10X commercial buffer. The mixture was incubated in room temperature for 1 hour. For full deglycosylation of receptors, 17 μl of the lysates was treated with 1 μl of Deglycosylation mix II (NEB, cat# P6044S) or distilled water, supplemented with 2 μl of their respective 10X commercial buffer. The mixture was first incubated in room temperature for 30 minutes, and then incubated in 37°C overnight. The reactions were terminated by adding Laemmli buffer. Samples were then analysed by SDS-PAGE and WB.

### Dynamic light scattering (DLS) studies

DLS measurements were performed using a DynaPro^®^ NanoStar^®^ instrument. Samples were centrifuged for 20 min at 21,130 g and 4°C, transferred into either a disposable plastic cuvette or a reusable quartz cuvette and allowed to equilibrate at 20°C in the instrument for 6 min before the measurement was started. Each measurement was averaged across 100 individual runs with an acquisition time of 5 s. Data for estimation of particle size of the different *h*B2 constructs (Table S2) were collected in a buffer consisting of 25 mM Tris, 150 mM NaCl, pH 8.0 (4°C) using a viscosity of 1.019 (20°C) and a refraction index of 1.333 (589 nm, 20°C).

### Crystal structure analysis

Diffraction quality crystals were obtained via hanging drop vapor diffusion by mixing 1 μL protein solution (5 mg/mL *h*B2-HG) with 1 μL reservoir solution, containing 100 mM sodium cacodylate pH 6.5, 200 mM MgCl2 and 20 % (w/v) PEG 8,000. Crystals grew as clusters of thin plates, which were separated by applying gentle pressure using a nylon loop. Crystals were flash frozen and stored in liquid nitrogen. Data was collected using synchrotron radiation at EMBL beamline P13 at the PETRA III storage ring (DESY, Hamburg, Germany), at λ = 1.77121 Å by an EIGER 16M detector. Crystals were kept at 100 K in a constant jet of nitrogen gas. XDS^65^ was used for indexing, integration and scaling of reflections. STARANISO^66^ was used to apply an anisotropic cutoff of merged intensity data. Final scaling and merging was carried out by AIMLESS^67^. Molecular replacement using the model 4dlo^4^ and refinement were carried out with the PHENIX^68^ suite while Coot^69^ was used for manual model building. Figures were generated using PyMOL (www.pymol.org). The PISA^70^ and EPPIC^71^ web servers were used to identify and characterize dimeric arrangements observed in the crystal structure. The refined model has been deposited in the Protein Data Bank as 8oek.

### Molecular dynamics simulations

#### ADGRL1 MD simulation

A model of the uncleaved L1 GAIN domain was constructed based on the crystal structure of the cleaved HormR-GAIN domain of rat ADGRL1 (PDB ID: 4dlq)^4^. The geometry of the peptide bond between the L^-1^ and T^+1^ residues of the GPS was manually modelled using the *Builder* utility in PyMol^72^. The CHARMM-GUI was used to normalize bond lengths and generate minimization and equilibration inputs using the CHARMM36 forcefield for GROMACS version 2020.2^73–76^. Default protonation states were verified by pKa analysis via Karlsberg2+^77^. The N-acetylglucosamine residues at N531, N640, N741, N800, N805, and N826 of the crystal structure were included in the model. Water boxes were generated with the CHARMM-GUI using a distance padding of 10 *Å* and charge-neutralized with 0.15 M NaCl. After minimization using the steepest-descent method for 5,000 steps, a 125,000 step equilibration with 1 fs timestep was performed to yield the equilibrated model. To obtain a starting structure that is competent for GPS hydrolysis, the unprotonated nitrogen atoms of the H^-1^ residues and the oxygen atom of the T^+1^ hydroxyl nucleophile were pulled together by a biased MD simulation. After 5 ns of equilibration with a time step of 1 fs, an harmonic umbrella potential was applied on both atoms with a force constant of 1000 kJ/mol/nm^2^ and a pull rate of −0.001 nm/ns until the run terminated due to a low-distance warning (HSD: 510 ps, HSE: 650 ps). With the resulting configurations, an equilibration cascade with decreasing harmonic potential holding the N_δ/ε_-O_γ_ distance constant for 100 ns with a decreasing force constant of 1000, 500, 200 and 100 kJ/mol/nm^2^, while simultaneously applying backbone, side-chain and dihedral restraints of 400, 400, 40 and 4 kJ/mol/nm^2^ or kJ/mol/deg, respectively, on the protein. After equilibration, triplicate unbiased MD simulations for 1,500 ns using the CHARMM36 force field in GROMACS were performed.

#### ADGRB2 MD simulation

The crystal structure of the B2 HrmR-GAIN domains determined in this work was used as a template for preparation of the MD model. The loop of residues 605-610, which was absent in the crystallographic model, was modeled using SuperLooper2^78^. The other, large missing section (residues 770-810) was deemed too large to be modeled and was left as a gap within the model. Buried waters within the structure were added using dowser^79^. MD system setup was carried out using CHARMM-GUI^73^, treating the termini of the introduced gap as methylated and acetylated and the protein C- and N-terminus as neutral. After evaluating residue pKa values using Karlsberg2+ with only H910 showing ambiguous pKa, three independent systems were setup with H910 as HSD, HSE and HSP, respectively. Glycosylations were added according to the UniprotKB entry O60241 at residues N560, N645 and N867. Water boxes were generated with CHARMM-GUI using a distance padding of 10 Å and charge-neutralized with 0.15 M NaCl. MD was performed using the CHARMM36 force field in GROMACS 2020.2. After minimization using the steepest-descent method for 5,000 steps, three independent 125,000 step equilibration runs with 1 fs timestep were performed per model before starting production runs with 2 fs timestep for 1,500 ns each.

MD analysis was carried out with GROMACS 2020.2 and MDtraj within a python3 environment and by using vmd^80^. Time-resolved pKa analysis was carried out via Karlsberg2+, calculating residue pKa for frames taken every 10 ns from each trajectory (151 frames per replica).

### Conservation analysis

For comparison of the different aGPCR amino acid sequences all available vertebrate sequences were extracted from NCBI and the respective orthologous sequences were separately aligned with program Muscle imbedded in the sequence analysis program Unipro UGENE v. 45.1^81^. Phylogenetic outliers (e.g. wrongly annotated sequences), sequences with missing sequence fragments, and misalignments were deleted from the alignments and not considered further. The conservation of the individual amino acid positions was determined with Unipro UGENE v. 45.1.

### Small angle X-ray scattering analysis

SAXS data was recorded at the EMBL SAXS beamline P12 at DESY ^82^ for a dilution series of *h*B2-HG between 0.5 mg/mL and 10 mg/mL (Fig. S20). The sample cell consists of a horizontal, temperature controlled (278-313 K) quartz cuvette, with a wall thickness of 50 μm and a path length of 1.7 mm. Data was collected at a wavelength of 1.23983 Å by a PILATUS 6M detector, positioned at a distance of 3 m. Approximately 30 μL of each sample continuously flowed through the quartz cuvette during the 0.1 s exposure window. These conditions were chosen in an effort to minimize radiation damage. The radius of gyration *R_g_* was determined by the Guinier approximation using PRIMUS, which is part of the data analysis suite ATSAS^83^.

In order to evaluate which species (monomer, dimer #1, dimer #3) best explains the experimental SAXS data, missing loop regions were added to the structure and 150 individual models were generated for each species, using MODELLER^84^. High-mannose type glycans were added to the predicted glycosylation sites Asn560, Asn646 and Asn867 in all models using the ALLOSMOD protocol^85^. Models with obvious clashes between sugar units and protein were excluded from further analyses, leaving a total of 118, 65 and 122 models for the monomer, dimer #1 and dimer #3, respectively. Differently glycosylated models were also tested, but yielded overall worse fits.

FoXS was then used to predict the scattering curves for individual models of the monomeric protein as well as potential dimers #1 and #3 and MultiFoXS was subsequently used to select ensembles out of the pool of the 305 individual species and refined the volume fraction of the models in the ensemble against the experimental data ^86^. The resulting ensembles contained between 2 and 5 models.

For these fits, summarized in table 3, the data up to *s* ≤ 3 nm^-1^ were included. A cut-off at small scattering vector length was determined for each sample and concentration based on a Guinier approximation using GNOM^87^. For estimation of the K_D_ value of the dimerization equilibrium, the volume fraction of the monomer as a function of protein concentration was used in a fit to the equation specified in Figure S20.

## Acknowledgements

We thank the Deutsche Forschungsgemeinschaft (DFG, German Research Foundation) for financial support through SFB1423, project number 421152132, subprojects A06 (T.L. and N.S.), B06 (T.L.), C01 and Z04 (P.H.), A05, C04 (T.S.). We acknowledge the EMBL beamlines of the DESY synchrotron in Hamburg for beamtime and support. We thank the MX Laboratory at the Helmholtz Zentrum Berlin (BESSY II) for beam time as well as for travel support.

## Author contributions

F.P.: Protein preparation, crystal structure analysis and SAXS studies; F.S.: MD simulations and analysis; Y.C.: GPS cleavage assays of full-length proteins; D.V.: MS analysis of furin cleavage; T.S.: conservation analysis; T.L., P.H., N.S.: Conceptualization of the studies; R.H., T.S., T.L., P.H., N.S.: supervision of the studies. F.P. wrote the first draft of the manuscript with contributions from F.S., Y.C. and D.V. All authors contributed to the interpretations and paper writing.

