## Supplemental Material for "Structural basis of GAIN domain autoproteolysis and cleavage-resistance in the adhesion G-protein coupled receptors"

-

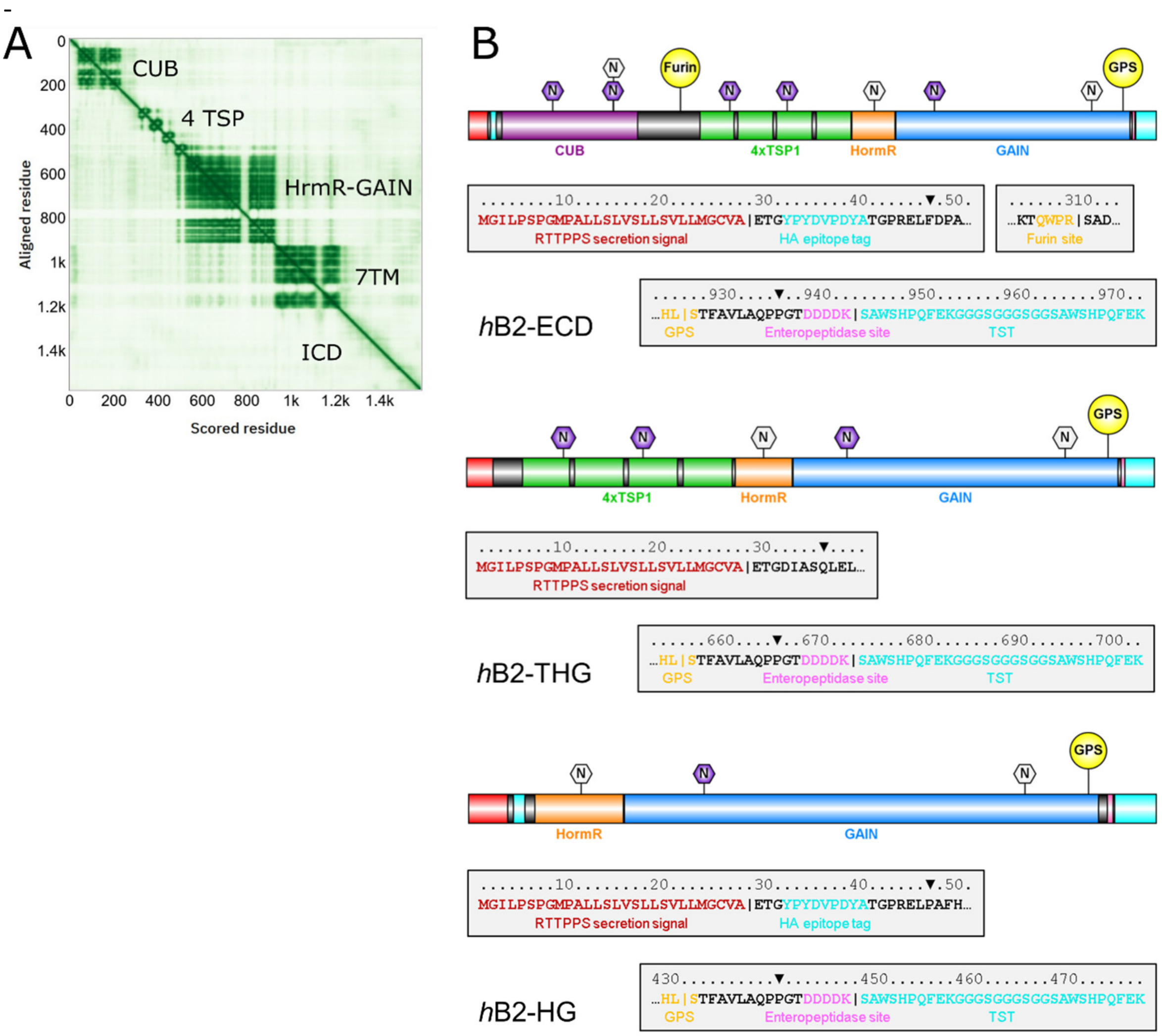


**Figure S1: AlphaFold prediction of the structure of ADGRB2 (A) and constructs including the GAIN domain used in this study (B).** (A) Predicted Aligned Error (PAE, green for small errors, white for large errors) of the AlphaFold2 prediction of the human B2 receptor. Low off-diagonal errors between domains are observed only for the HormR and GAIN domains, and to a lesser extent between the last thrombospondin (TSP) domain and the HormR-GAIN domains. This indicates that these regions are likely to have interactions and have a stable orientation relative to each other. In contrast, all other domains are predicted not to interact. Longer disordered (flexible) regions are predicted between the CUB and the first TSP domain and for the whole intracellular region. (B) Expression constructs *h*B2-ECR (top), *h*B2-THG (middle) and *h*B2-HG (bottom). The secretion signal is cleaved off before the mature protein is secreted into the medium. Possible N-glycosylation sites (N) were predicted by NetNGlyc 1.0. Those with a potential greater than 0.5 are shaded in purple. CUB – complement C1r/C1s, Uegf, Bmp1 (domain); GAIN – GPCR autoproteolysis inducing (domain); GPS – GPCR proteolytic site; HA – human influenza hemagglutinin; HormR – hormone receptor (domain); RTTPPS – receptor-type tyrosine-protein phosphatase S; TSP1 – thrombospondin-type 1 (repeat); TST – Twin-Strep-tag®; | denotes expected cleavage sites; ▼ denotes the insert's boundaries.

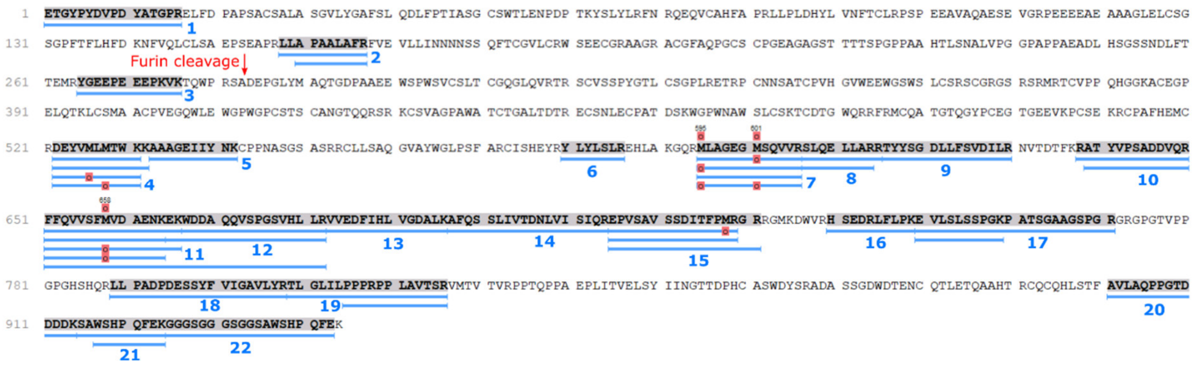


**Figure S2: Peptides detected in the mass spectrometric protein identification of the bands shown in Figure 2**. Peptides 1-3 and 5 have been used to demonstrate the absence of peptides 1-3 in the band corresponding to the dimer after furin cleavage. Thus, this dimer is a homodimer and no heterodimers with one furin-cleaved subunit are observed. Peptide 5 has been used as a control, this peptide is located downstream of was observed

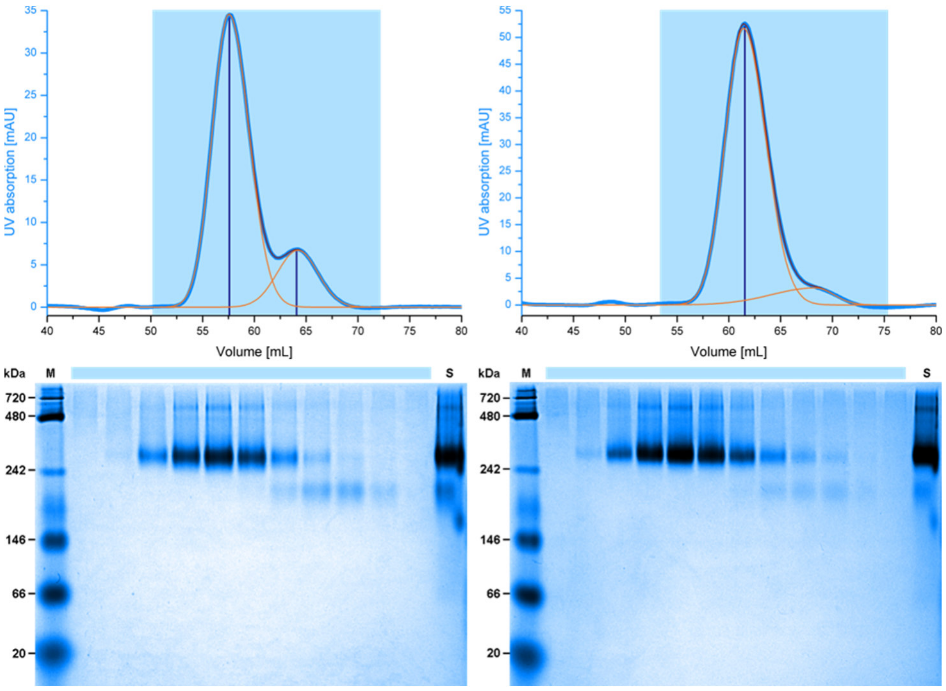


**Figure S3: Reduction of furin cleavage in the Q292A/R295A double mutant of the furin recognition site.** Comparison of *h*B2-ECR (left) and the Q292A/R295A double mutant *h*B2-ECR-DM (right) SEC purification profiles (top). Baseline correction was applied to the size exclusion chromatography (SEC) profiles (top) before Gaussian curves with individual slopes for the front- and backside (to account for asymmetric elution peaks) were fit to the data in order to estimate the amount of furin-cleaved protein. The cleaved fraction was reduced from 15.6 % in the wild type to 7.4 % in the double mutant and clearly still visible in a BN-PAGE (bottom). M – marker; S – sample before application to SEC.

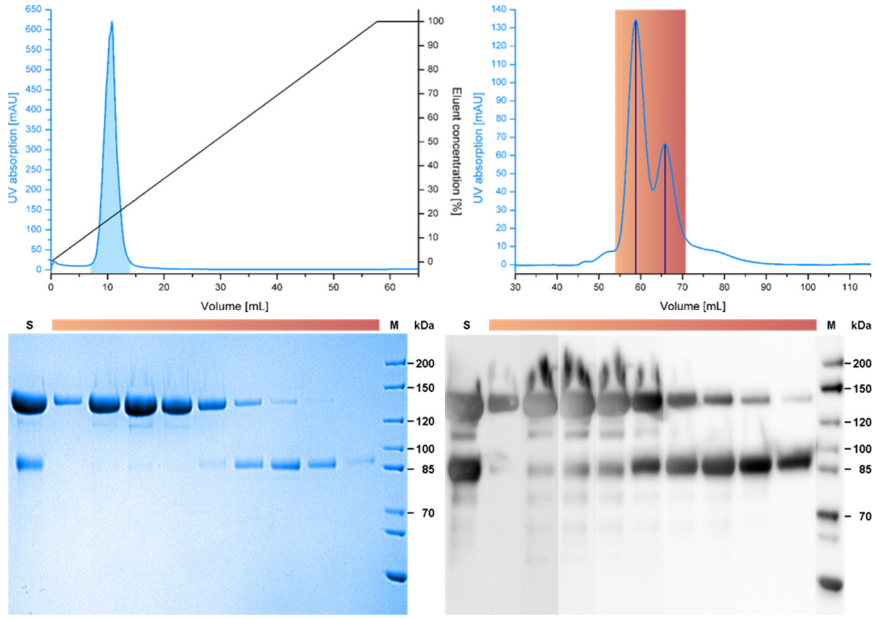


**Figure S4: Purification of hB2-ECR**. The chromatograms of the affinity chromatography (top left) and SEC (top right) as well as an SDS-PAGE under nonreducing conditions (bottom left) and subsequent western blot detected via the C-terminal TST (bottom right) of the SEC illustrate the purification progress. Two distinct but overlapping peaks were observed in the SEC, which were attributed to two homodimer species of the full-length protein (larger peak, upper band) and the C-terminal fragment following furin cleavage (smaller peak, lower band). Some degradation of the protein was visible. M – marker; S – sample applied to the SEC. Theoretical and determined masses are listed in Table 1. The western blot shows that the second peak of the SEC profile contains a homodimer of two furin-cleaved subunits as the peak maximum does not contain equal amounts of the two chains.

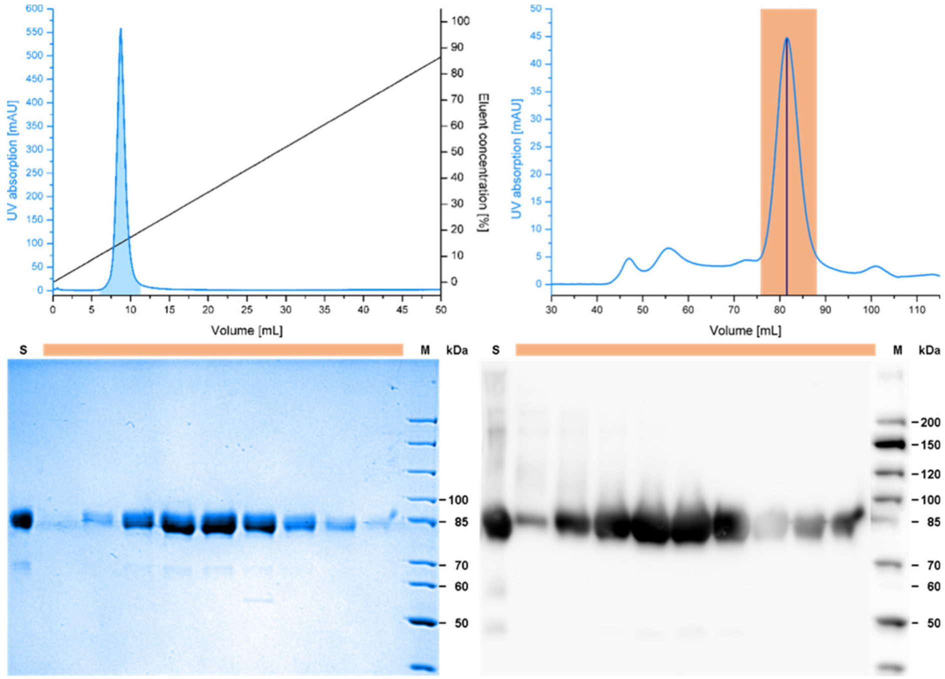


**Figure S5: Purification of hB2-THG**. The chromatograms of the AC (top left) and SEC (top right) as well as an SDS-PAGE under nonreducing conditions (bottom left) and subsequent western blot detected via the C-terminal TST (bottom right) of the SEC illustrate the purification progress. The fractions marked in orange in the SEC have been analyzed by SDS-PAGE. Some degradation of the protein was visible as faint bands in the SDS-PAGE and western blot. The western blot showed a faint smear of likely oligomerized protein present in the earlier peak fractions. M – marker; S – sample applied to the SEC. Theoretical and determined masses are listed in Table 1.

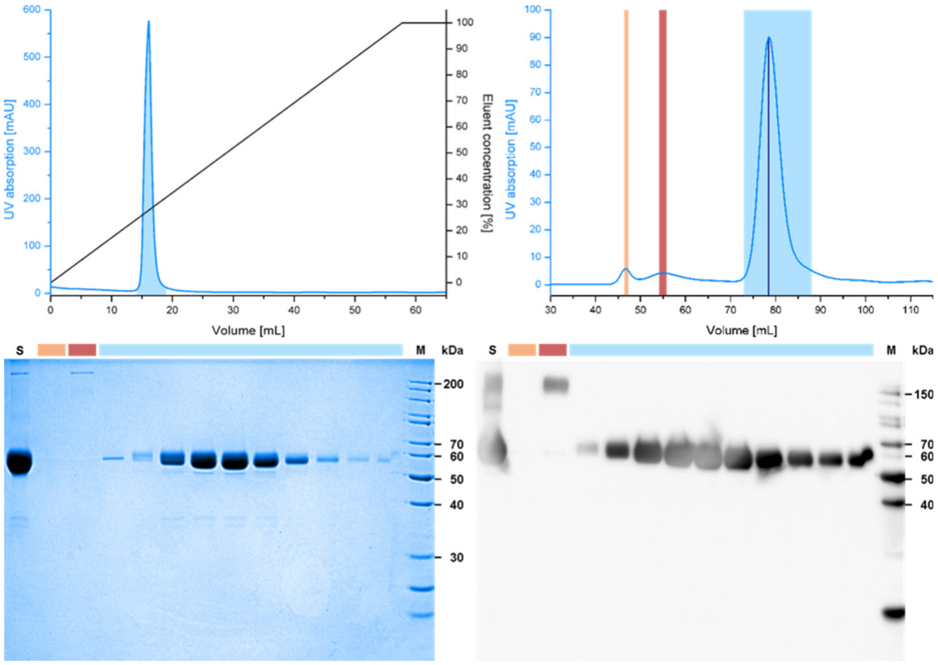


**Figure S6:** Purification of hB2-HG. The chromatograms of the AC (top left) and SEC (top right) as well as an SDS-PAGE under nonreducing conditions (bottom left) and subsequent western blot detected via the C-terminal TST (bottom right) of the SEC illustrate the purification progress. A small amount of cysteine-bridged oligomer (orange) was separated from the main peak (blue), which exhibited minor tailing in the SEC. Some degradation of the protein was visible as faint bands in the SDS-PAGE. M – marker; S – sample applied to the SEC. Theoretical and determined masses are listed in Table 1.

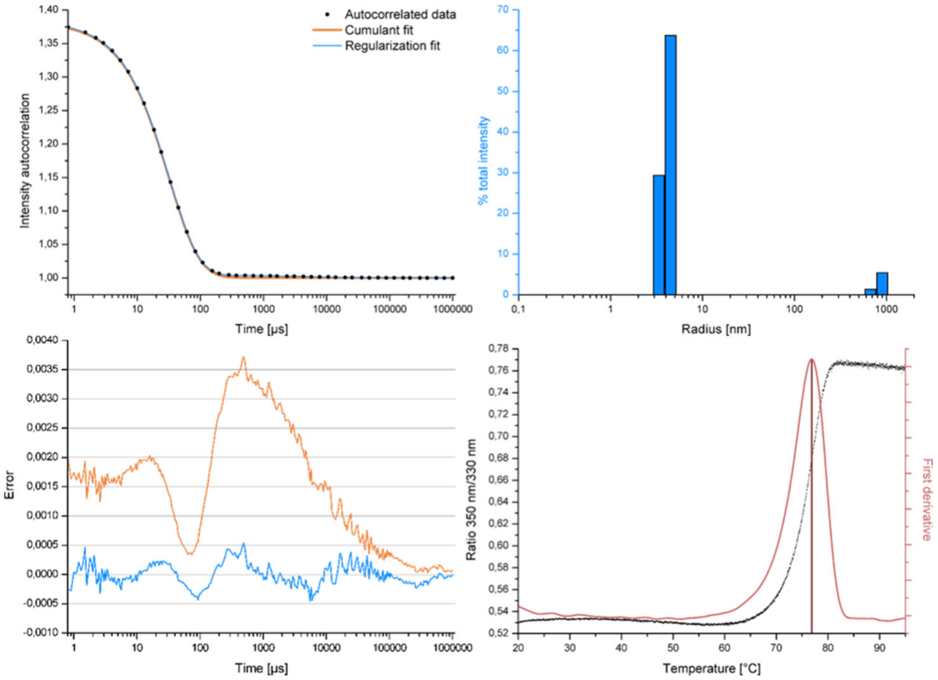


**Figure S7: Dynamic light scattering (DLS) and differential scanning fluorimetry (DSF) analysis of purified hB2-HG**. The DLS autocorrelated data with cumulant and regularization fits is shown on the top left and the corresponding errors on the bottom left. The results of the regularization fit are visualized as a bar diagram on the top right. The DLS, measured at 10 mg/ml, showed a main species with a radius of ~4.1 nm (94 kDa for a spherical protein) and a polydispersity of 12.9 % with minor contamination by a larger, likely aggregated species. The DSF on the bottom right showed a sharp transition with a melting point of 76.6±0.08°C (n=3).

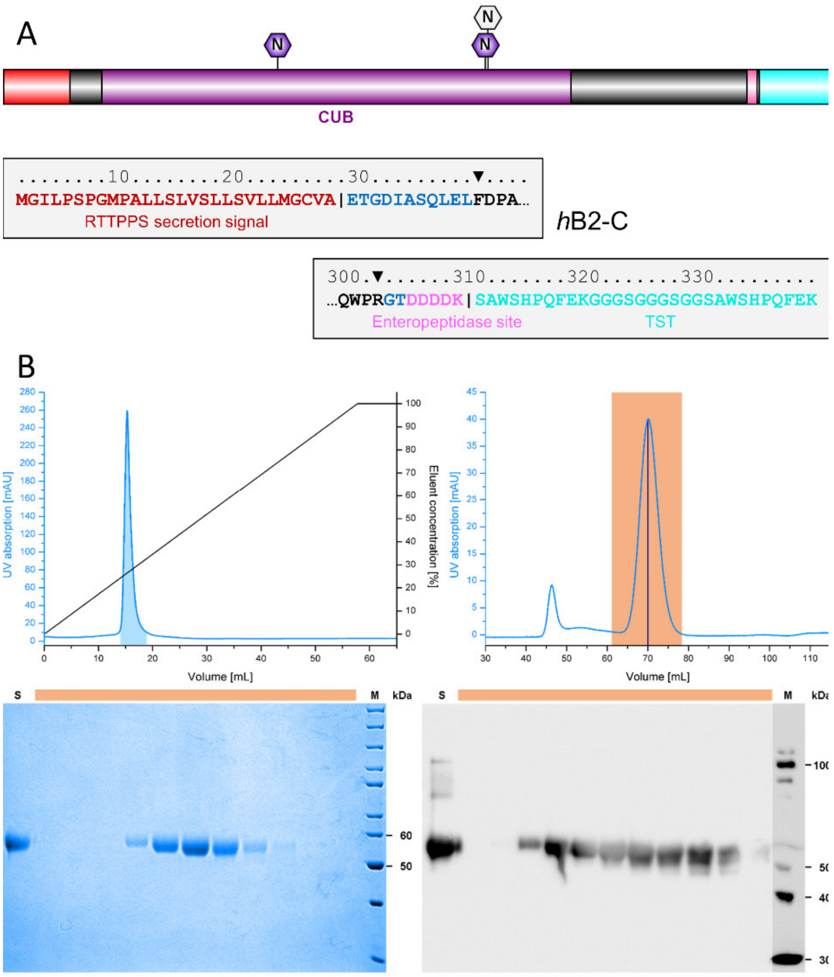


**Figure S8: Purification of *h*B2-C.** (A) Expression construct hB2-C. The secretion signal is cleaved off before the mature protein is secreted into the medium. Possible N-glycosylation sites (N) were predicted by NetNGlyc 1.0. Those with a potential greater than 0.5 are shaded in purple. CUB – complement C1r/C1s, Uegf, Bmp1 (domain); RTTPPS – receptor-type tyrosine-protein phosphatase S; TST – Twin-Strep-tag®; | denotes expected cleavage sites; ▼ denotes the insert's boundaries. The sequence regions shown in blue are derived from the expression vector. (B) Chromatograms of the AC (top left) and SEC (top right) as well as an SDS-PAGE under reducing conditions (bottom left) and subsequent western blot detected via the C-terminal TST (bottom right) of the SEC illustrate the purification progress. The peak of the desired protein (orange) gave rise to a symmetrical peak in the SEC. M – marker; S – sample applied to the SEC. Theoretical and determined masses are listed in Table 1.

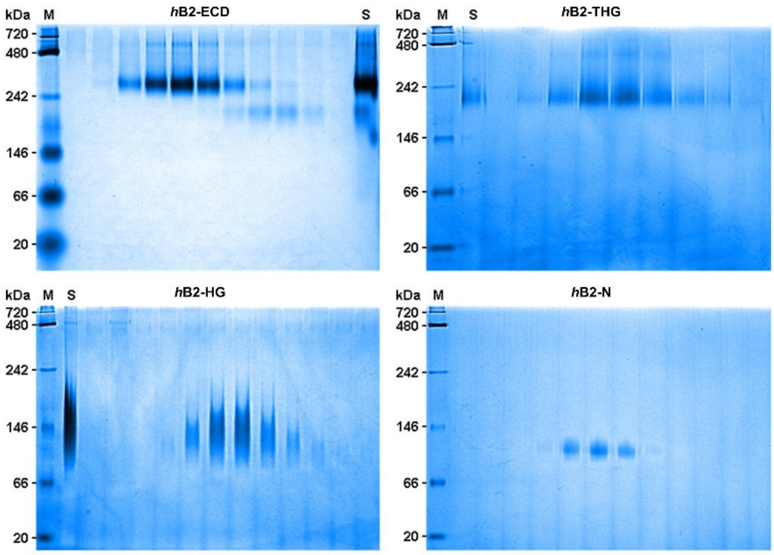


**Figure S9:** BN-PAGE analysis of the SEC fractions of *h*B2-ECR, *h*B2-THG, *h*B2-HG and *h*B2-C. M – marker; S – sample applied to SEC. Theoretical and determined masses are listed in Table 1.

**Table S1:** Hydrodynamic radii ($R_h$) and molecular weights (MW) estimated by DLS. It should be noted that the molecular weights calculated from the radius of hydration ($R_h$) determined by the DLS experiment assume a spherical protein shape and are strongly dependent on a correct estimation or determination of the viscosity of the buffer. The statistical standard deviation given for these values thus highly overestimates the accuracy of these measurements.

| | ***h*B2-ECR *(n=6)*** | ***h*B2-THG *(n=2)*** | ***h*B2-HG *(n=8)*** |
|---|---|---|---|
| c [mg/ml] | 1.0 | 1.0 | 5.0 |
| $R_h$ [nm] | 7.285 ± 0.063 | 5.853 ± 0.017 | 4.150 ± 0.029 |
| MW [kDa] | 351.0 ± 7.0 | 210.5 ± 1.5 | 94.1 ± 1.4 |
| theoretical mass [kDa] | | | |
| monomer | 108.6 | 79.7 | 55.4 |
| dimer | 217.2 | 159.4 | 110.8 |

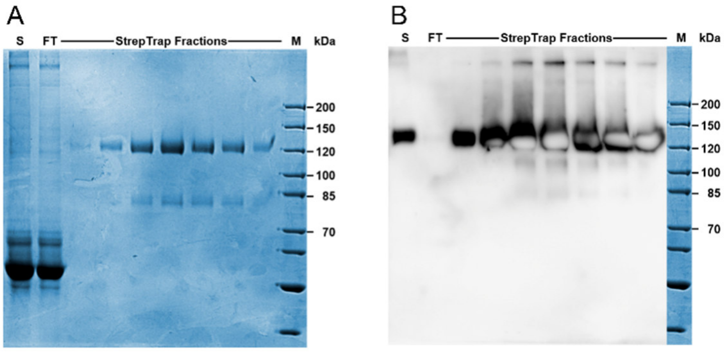


**Figure S10: Furin cleavage results in symmetrically cleaved homodimers.** (A) Coomassie-stained non-reducing SDS-PAGE of the fractions of the main peak eluted from the initial Strep-Tactin affinity chromatography. Bands are observed at apparent molecular masses of 120 and 81 kDa. (B) A western blot against the N-terminal HA-tag shows that only the band observed at 120 kDa carries a HA-tag. Thus, the band observed at 81 kDa is a symmetrical dimer obtained after cleavage of the furin cleavage sites in both chains. No heterodimer is observed, in which only one chain is processed, in agreement with the results shown in Figure 2. In addition, this analysis indicates that no partial GPS clavage occurs as no double bands are observed, which would be expected from a removal of the *Stachel* peptide (theoretical mass 4.8 kDa) after GPS autoproteolysis. Expected molecular masses are specified in table 1.

**Table S2: Crystallization of hB2-HG.** Protein samples were used in the respective SEC buffers and their concentration is listed below. Crystals usually appeared within 48 h and kept slowly growing over time. Images shown here were taken after 34 days. Crystals appeared in ~12 out of ~700 tested conditions in the initial screening. The needle-like or plate-like crystals always grew from a common nucleation point as clusters.

| Protein concentration | Crystallization condition | Crystals |
|---|---|---|
| *h*B2-HG<br>5 mg/ml | **M1B6**<br>100 mM sodium cacodylate, pH 6.5<br>200 mM magnesium acetate<br>20 % (w/v) PEG 8,000 | 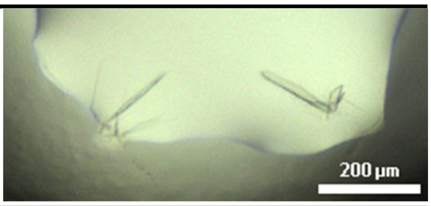  |
| *h*B2-HG,<br>deglycosylated<br>5 mg/ml | **M1B6**<br>100 mM sodium cacodylate, pH 6.5<br>200 mM magnesium acetate<br>20 % (w/v) PEG 8,000 | 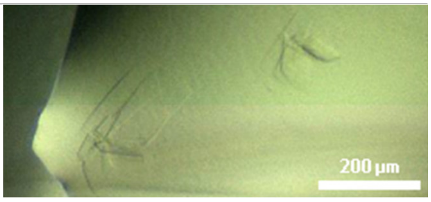  |
| *h*B2-HG<br>5 mg/ml | **M4F3**<br>100 mM MES, pH 6.0<br>20 % (w/v) PEG 6,000 | 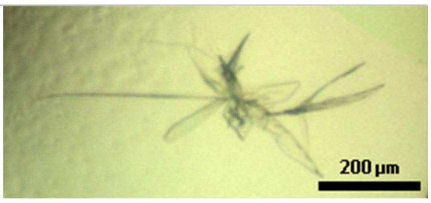  |

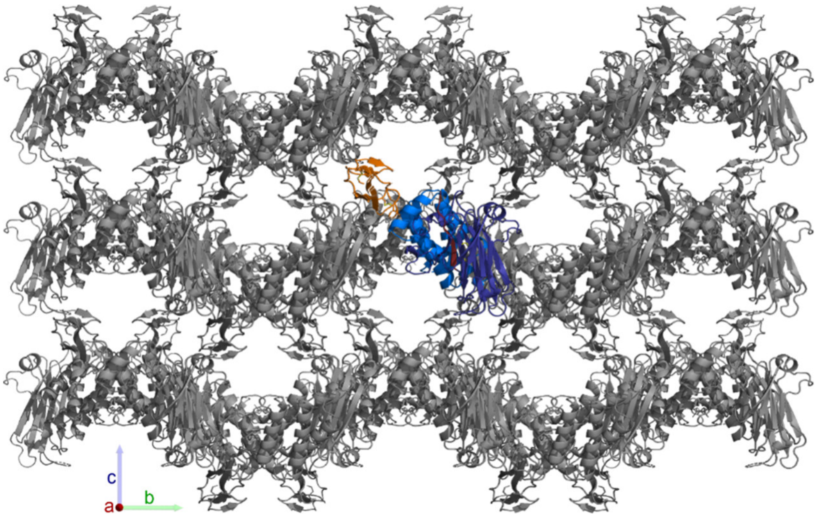


**Figure S11:** Crystal packing of *h*B2-HG in space group $P2_12_12$. Crystals grew in layers with large channels between them. Crystal contacts between the layers are formed by the HormR domains. The axes shown on the bottom left align with the crystallographic unit cell. The weak contacts along the direction of unit cell axis **c** likely cause the platelike growth of the crystals (the **c**-axis located normal to the plates) and the anisotropic diffraction limits of 2.21 Å, 2.31 Å , and 3.02 Å along the **a***, **b***, and **c*** axes, respectively, observed for the crystal structure. The monomer is shown with the HormR domain in orange and the GAIN domain in blue.

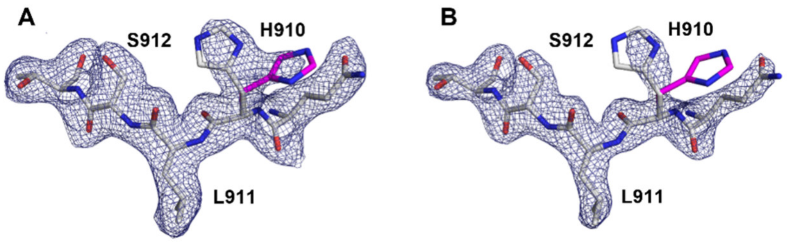


**Figure S12: Conformational flexibility of the H910 side chain in the *h*B2-HG crystal structure.** The sidechain of H910 is present in two distinct orientations. The electron density around the GPS is shown at a contour level of 1.3 σ (A) and 0.6 σ (B). The two alternative rotamers of the H910 sidechain are shown in grey (refined occupancy 0.57) and magenta (0.43).

**Table S3: Summary of crystallization, data collection and refinement**

| | ***h*B2-HG (pdb ID 8oek)** |
|---|---|
| **Final crystal buffer** | 100 mM sodium cacodylate, pH 6.7<br>200 mM magnesium acetate<br>18 % (w/v) PEG 8,000 |
| **Data collection** | |
| Wavelength [Å] | 1.77121 |
| Resolution limits [Å] | 63.34 - 2.22 (2.45 - 2.22) |
| Diffraction limits a*, b*, c* [Å] | 2.21, 2.31, 3.02 |
| Space group | $P2_12_12$ |
| Unit cell a, b, c [Å] | 84.86, 95.18, 49.46 |
| Total reflections | 164790 (6912) |
| Unique reflections | 13145 (658) |
| Multiplicity | 12.5 (10.5) |
| Completeness (spherical) [%] | 63.8 (12.6) |
| Completeness (ellipsoidal [%] | 89.1 (48.6) |
| Mean I/σ(I) | 9.9 (1.3) |
| $R_{meas}$ | 0.223 (1.949) |
| $R_{pim.}$ | 0.062 (0.583) |
| CC½ | 0.997 (0.567) |
| Wilson B-factor [$Å^2$] | 29.07 |
| **Refinement** | |
| Resolution range [Å] | 42.75 - 2.22 (2.30 - 2.22) |
| $R_{work}$ | 0.2325 (0.3803) |
| $R_{free}$ | 0.2869 (0.3499) |
| Number of non-hydrogen atoms | |
| *Protein* | 2717 |
| *Heterogen* | 33 |
| *Solvent* | 18 |
| B-factors [$Å^2$] | |
| *Protein* | 35.39 |
| *Heterogen* | 49.20 |

| | |
|---|---|
| *Solvent* | 32.23 |
| Ramachandran statistics [%] | |
| *Favored* | 96.45 |
| *Allowed* | 3.25 |
| *Outliers* | 0.30 |
| Root mean square deviation (RMSD) | |
| *Bond lengths* [Å] | 0.002 |
| *Bond angles* [°] | 0.43 |

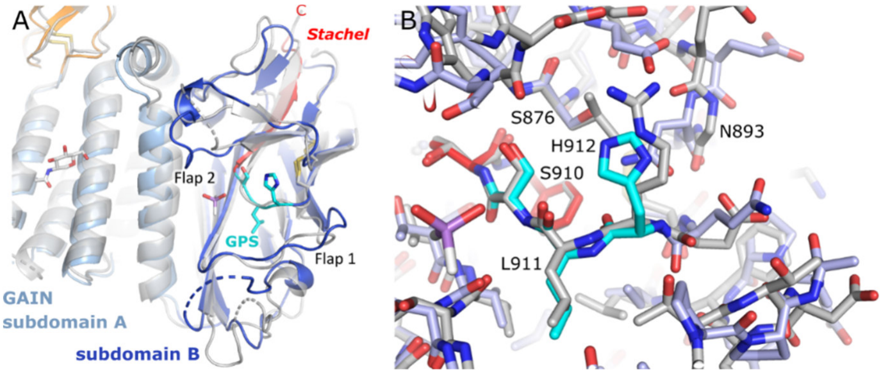


**Figure S13: Comparison of the GPS environment in the B2 and B2 GAIN domains.** (A) Superposition of the crystal structures of the GAIN domains of *h*B2 (colors) and *h*B3 (grey, pdb id 4dlo). The two structures differ significantly in the fold of the flap regions around the GPS. (B) Closer view of the immediate GPS environment (B2 in colors, B3 in grey). Noteworthy differences are N893, which corresponds to the highly conserved glycine in B3 and in cleavage-competent GAIN domains, and S876, which corresponds to the phenylalanine or tyrosine residue forming an edge-π interaction with the histidine base (H911 in *h*B2) in the cleavage-competent receptors and is a leucine in *h*B3. Furthermore, the histidine base is replaced by an arginine residue in the GPS motif of *h*B3. Many other residues in the GPS environment also differ in sequence or conformation. The structures have been superimposed based on subdomain B of the GAIN domain.

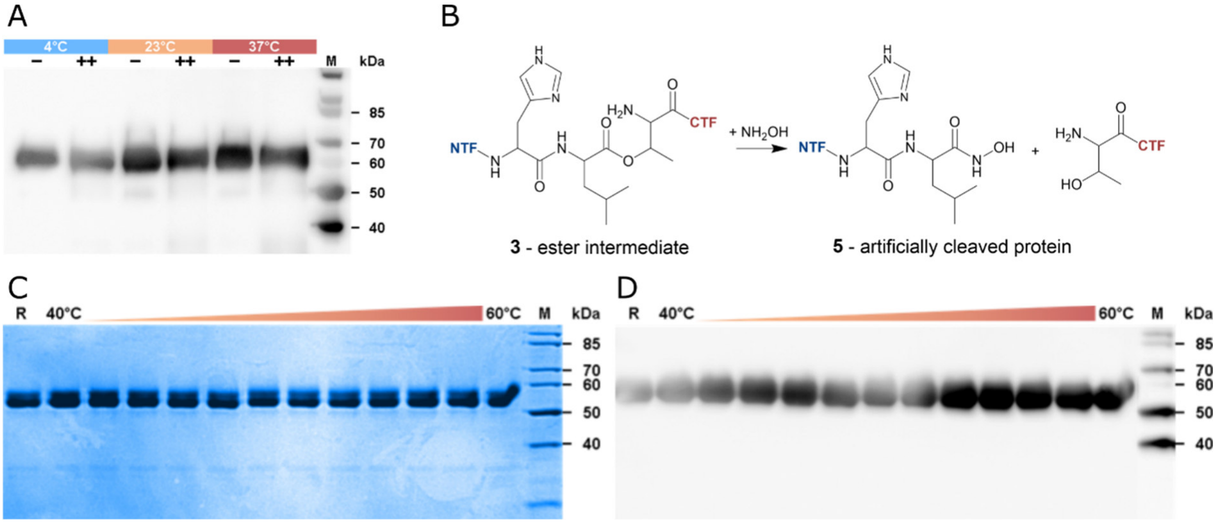


**Figure S14: The B2 GAIN domain is resistant to cleavage at the GPS even under elevated temperatures or treatment with hydroxylamine.** (A) Hydroxylamine assay of *h*B2-HG. – no hydroxylamine added; + 250 mM hydroxylamine; ++ 500 mM hydroxylamine; M – marker. The samples were incubated for overnight at the specified temperatures in a buffer containing 25 mM Tris pH 8.0 (determined at 4°C) and 150 mM NaCl prior to SDS-PAGE analysis. (B) Scheme of the hydrolysis reaction of an ester intermediate by hydroxylamine (C) Heat treatment of purified *h*B2-HG. The protein was incubated for 1 h at 40-60°C in a buffer containing 25 mM Tris pH 8.0 and 150 mM NaCl before the samples analyzed via SDS-PAGE under reducing conditions. (D) Western blot of the SDS-PAGE gel detected via the C-terminal TST. No shift in the apparent molecular weight or vanishing of bands in the western blot were observed.

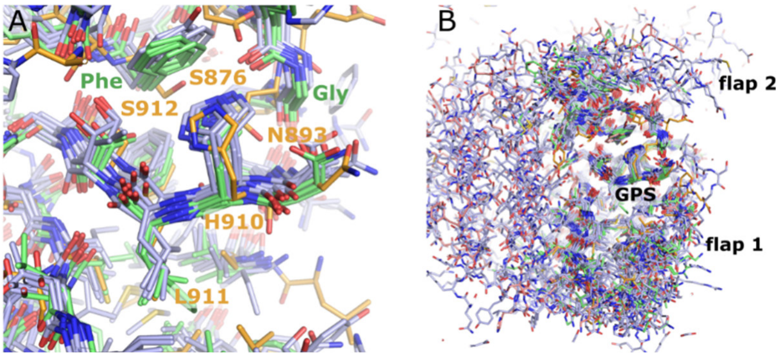


**Figure S15: Comparison of the GPS and its environment between the B2 GAIN domain and cleavage-competent receptors.** (A) Comparison of the GPS and its environment between AlphaFold models of the following human receptors. Shown are cleavage-competent aGPCR for which structures of cleaved GAIN domains have been determined (green, L1, G1, G3 and G6) and for which cleavage has been reported mostly via gel-electrophoresis experiments: C2, D1, E2, E3, E5, F1, F3, F5, G2, G4, and L4 (blue). The crystallographic model of B2 is shown in orange. The histidine base of the GPS motif is oriented by an edge-π interaction with a phenylalanine (or tyrosine in E3) in cleavage-competent receptors whereas this residue is S912 in *h*B2. Furthermore, a highly conserved glycine residue is observed in autoproteolysis-active receptors at the position of N893 in *h*B2. (B) The wider GPS environment in these models differs tremendously due to the diversity and conformational flexibility of the flap regions and neighboring loops.

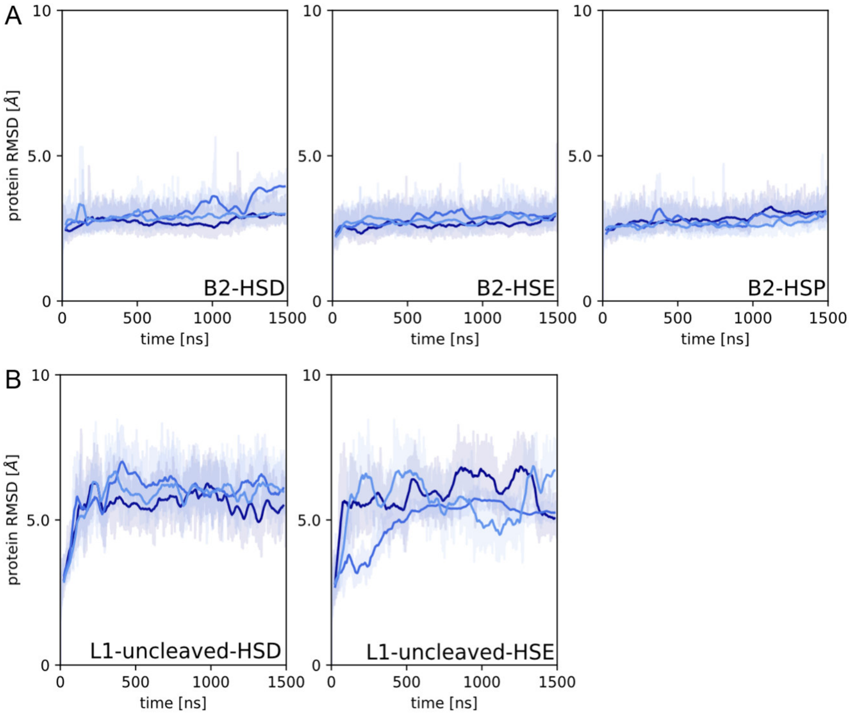


**Figure S16: Development of root mean square deviations (RMSD) to the starting structure during the time course of the MD simulations.** (A) RMSD plots for the three replicas (dark blue: r1, medium blue: r2, bright blue: r3) of the three simulations (HSD, HSE and HSE) of the B2 GAIN domain. (B) RMSD plots for the HSD and HSE simulations of the L1 GAIN domain. The colored RMSD traces are averaged over a window of 20 frames, whereas the plots in grey indicate the RMSD fluctuations for each frame.

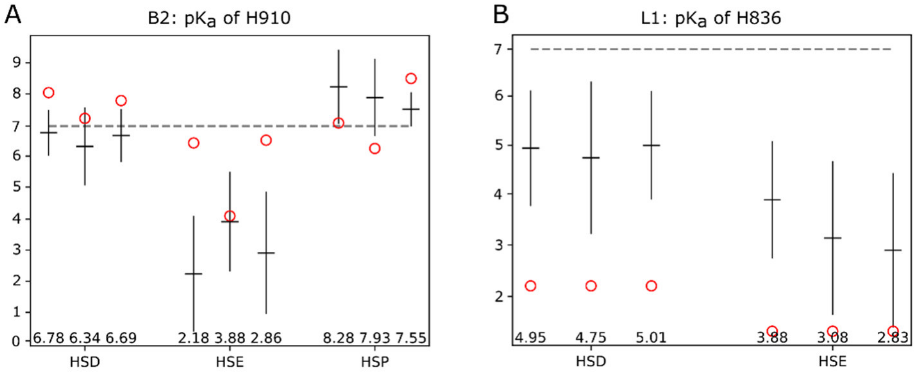


**Figure S17: Prediction of $pK_a$ values for the catalytic histidine bases of the GPS during B2 and L1 MD simulations.** (A) The pKa value if H910 was predicted by solving the Poisson−Boltzmann equation for each frame of the three replica of the HSD ($N_\delta$ protonated), HSE ($N_\varepsilon$ protonated) and HSP (both nitrogens protonated) protonation states. The average $pK_a$ values are indicated by the horizontal line and specified numerically. The range of pKa values are depicted by the vertical lines. Red circles mark the pKa value of the starting structure after equilibration (B) $pK_a$ values of H836 of the L1 simulations.. As the HSE state results in MD conformations with the lowest $pK_a$ values, the HSE protonation state is most stable at pH 7.

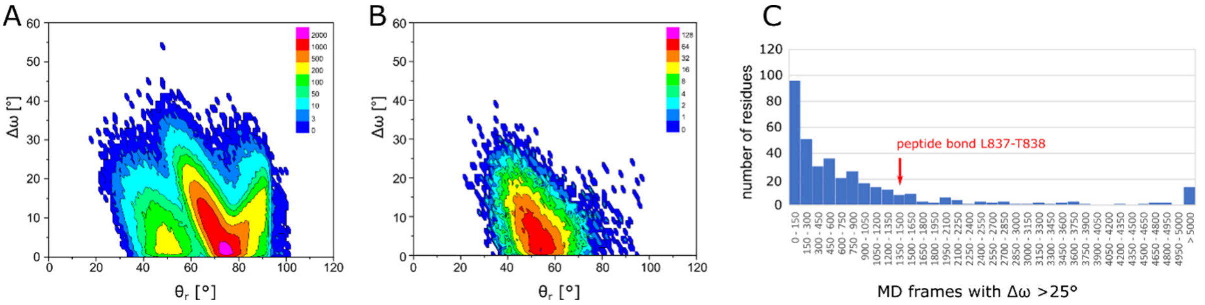


**Figure S18: Strain on the scissile peptide bond in the MD simulation of L1-HSE.** (A) Scatterplot of $\Delta\omega$ (the deviation of the peptide bond from planarity) and $\Theta_r$ (the angular deviation from the perfect $\Theta_x = 90°$ and $\Theta_y = 90°$ orientation for attack of the alcohol nucleophile on the peptide bond). $\Delta\omega$ and $\Theta_r$ was determined for all frames of the 3 replicas of the L1-HSE simulation. The scales indicate the color coding to visualize the number of frames for which a given pair of $\Delta\omega$ and $\Theta_r$ (binned in 1° steps) is observed. The three maxima correspond to different rotamers of the T838 nucleophile. As only the p-rotamers of T838 result in reactive conformations of the GPS, frames with this rotamer of T838 (criterium $\chi_1 = 59 \pm 30°$) are included in the scatter plot in (B). With decreasing deviation from the ideal $\Theta$ angles for nucleophilic attack, the strain $\Delta\omega$ of the scissile peptide increases. (C) Histogram showing how often residues of the HormR GAIN domains of L1 show strained peptide bonds ($\Delta\omega >$ 25°) in replica 3 of the L1-HSE simulation. For example, around 85 residues have $\Delta\omega$ angles larger than 25° in less than 150 of the 150706 frames of this simulation. Around 15 residues have strained peptide bonds in more than 5000 frames. For the scissile peptide bond L837-T838 $\Delta\omega > 25°$ was observed in 1550-1650 frames. Thus, strained conformations occur for the scissile peptide bond more often than for most of the other residues (82.7 %).

**Table S4: Geometric parameters describing the orientation of the S/T alcohol nucleophile, the histidine base and the scissile peptide bond in the HSD ($N_\delta$ is protonated) MD simulations of the B2 and L1 GAIN domains**. The values specify the number or percentage of frames in which the given condition is satisfied. These simulations were carried out with a neutral histidine base protonated at $N_\delta$. Results from simulations with protonation at $N_\varepsilon$ are shown in Table 2.

| | **$\Theta_x = 90° \pm 30°$ and $\Theta_y = 90° \pm 30°$** | **Hbond*** | **Hbond and $\Theta_{x/y} = 90° \pm 30°$** | **$\chi_1$** His +60°, -60°, -180°** | **$\chi_1$** Ser/Thr +60°, -60°, -180°** |
|---|---|---|---|---|---|
| **B2 r1** | 952 | 20 | 0 | 45.2%, 39.1%, 9.2% | 7.2%, 9.8%, 74.3% |
| **B2 r2** | 658 | 20 | 0 | 33.9%, 48.5%, 11.2% | 7.7%, 8.5%, 73.9% |
| **B2 r3** | 1054 | 18 | 0 | 47.9%, 36.7%, 8.6% | 11.0%, 13.4%, 65.9% |
| **L1 r1** | 3044 | 22 | 1 | 6.8%, 78.0%, 7.8% | 42.0%, 1.1%, 24.7% |
| **L1 r2** | 2109 | 9 | 0 | 10.6%, 66.4%, 15.6% | 41.8%, 0.3%, 28.8% |
| **L1 r3** | 27 | 0 | 0 | 5.1%, 72.4%, 10.2% | 11.5%, 0.6%, 52.8% |

*A favorable hydrogen bonding geometry was defined as a distance of 3.2 Å or less between the histidine nitrogen atom and the Ser/Thr $O_\gamma$ atom and a deviation of less than 30° of the $O_\gamma$-H···N angle from linearity.

**The $\chi_1$ torsion angle was assigned to the given value if it deviated by less than 30° from this value.

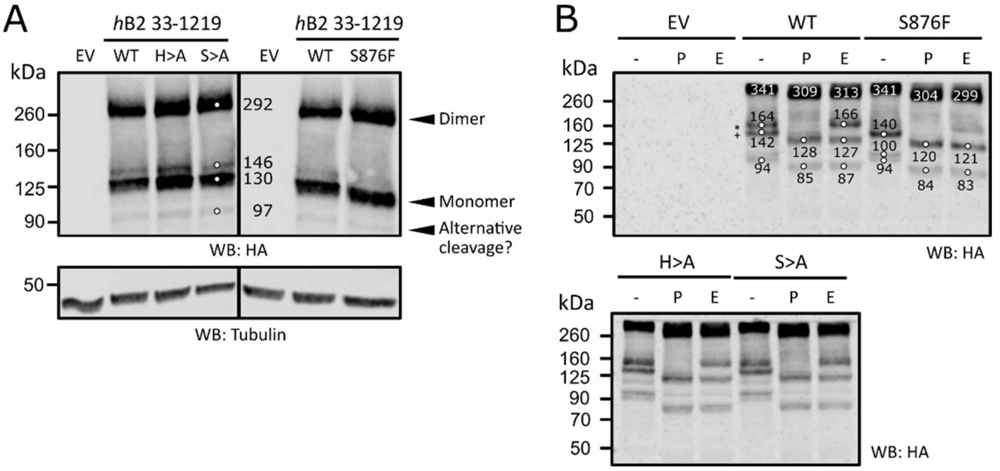


**Figure S19: Autoproteolysis is not observed for the full-length *h*B2 (further details).** HEK293T cells were transiently transfected with indicated mutated constructs of *h*B2 residues 33-1219. (A) Western blotting of an SDS-PAGE showed the expressions of *h*B2 and respective mutants targeted against the N-terminal HA tag of the receptor. Thirty µl of lysate samples were used for analyses. Tubulin served as loading control. For the bands indicated by the small open circles, the apparent molecular mass has been estimated using the software GelAnalyzer 19.1 (www.gelanalyzer.com) to the values (in kDa) indicated next to the bands. (B) Western blotting of SDS-PAGE of *h*B2 lysates treated with PNGase F ("P"), Endo H ("E"), or without any enzyme treatment (-). Endo H-resistant and sensitive bands of monomeric *h*B2 indicated by "*" and "+", respectively. The ECD of B2 contains five probable and three further N-glycosylation sites of lower probability (Fig. S1B). The calculated molecular masses of the used construct are 137.6 kDa (157.6 kDa with 8 N-linked complex glycans of 2.5 kDa) for full-length *h*B2, 32.8 kDa (40.3 kDa with 3 N-glycans) for the N-terminal fragment up to the furin cleavage site, 100.4 kDa (120.4 kDa with 8 N-glycans) for the NTF. We interprete the bands observed at ~130 kDa and ~290 kDa as uncleaved full-length receptor migrating as monomer and dimer. Membrane proteins can display significant anomalies in migration behavior in SDS-PAGE[1]. O-glycosylation likely contributes to further deviations from the expected molecular mass.

The band marked as "Alternative cleavage?" is observed in all samples at an apparent molecular mass of ~90 kDa. This band is also observed in the GPS mutants H910A and S912A, which are highly unlikely to possess autoproteolytic activity. These fragments may result in cleavage of the long disordered loop 770-810, which would generate a fragment of calculated mass of 87.2 kDa (104.7 kDa for 7 N-glycans) for cleavage at position 790. PNGase F removes the complete glycan chain for all types of N-linked glycosylation (high mannose, hybrid, bi-, tri-, and tetra-antennary), unless the innermost GlcNAc residue is α1-3-linked to a fucose. Endo H removes only high mannose and some hybrid types of N-linked carbohydrates by cleavage between the first and second GlcNAc residues. The two bands observed at apparent molecular masses of 164 kDa and 142 kDa for the deglycosylation experiment in panel (B) differ therefor in N-glycosylation. Band "*" contains additional complex glycans that are not removed by EndoH. Band "+" contains only glycans that can be removed by Endo H. It is interesting to note that predominantly only two distinct bands are observed. These differ in the maturation state of several glycans such that all are present as complex glycans in "*" and are missing or present in a state that the can be processed by Endo H in "+".

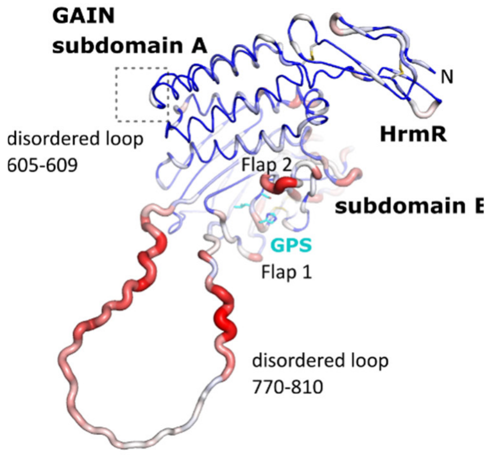


**Figure S20: Conservation of the long flexible loop 770-810 of B2.** Sequences of the HormR and GAIN domains of 148 mammalian species were aligned and the Shannon entropy was calculated for each residue, which quantifies the variability of the sequence positions. Residues of high conservation are shown as blue thin tube regions and residues of high sequence variation as red thick tube regions. This figure includes the loops of residues 605-609 and 770-810, which are disordered in the crystal structure. The loops are modeled in arbitrary conformations only for visualization of the residue conservation.

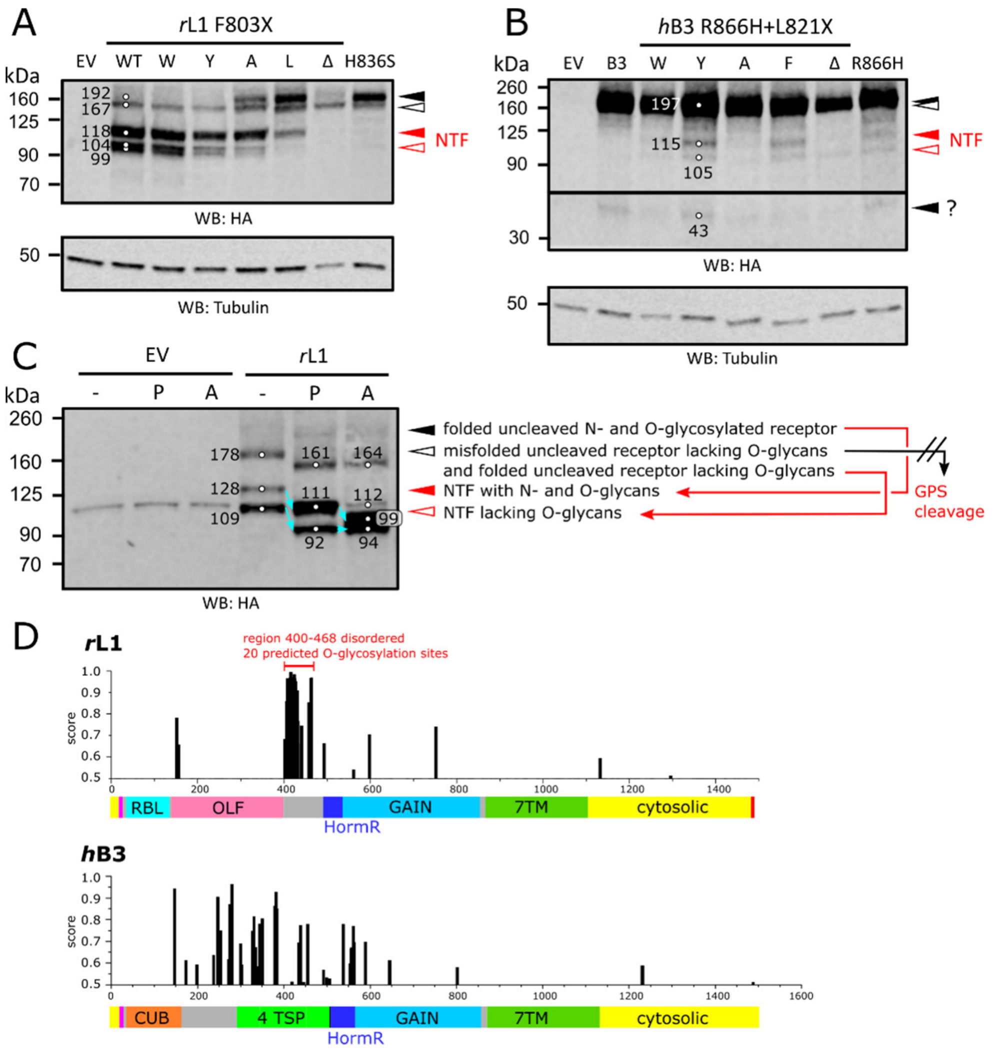


**Figure S21: Edge-π interaction between His$^{-2}$ base and surrounding residues is crucial for GPS cleavage of some receptors (further details and interpretation).** (A,B) HEK293T cells were transiently transfected with constructs containing mutations causing either a disruption of edge-π interaction in rL1 (A), or a reintroduction of edge-π interaction in *h*B3 (B). GPS cleavages of the mutants were analyzed by WB using 30 µl of cell lysates, targeted against the N-terminal HA tag of the receptors. Tubulin served as a loading control. Bands representing uncleaved and cleaved populations were indicated in black and red triangles, respectively. Fully and immaturely glycosylated populations of rat L1 were indicated in closed and open triangles, respectively. (C) rL1 is N- and O-glycosylated. HEK293T cells were transiently transfected with rL1. Lysates were treated with PNGase F ('P'), Deglycosylation mix II ('A') or without enzyme ('-'). Glycosylation state of receptors were examined by mobility shift of bands between treated and untreated samples (red dotted lines), detected against the N-terminal HA tag of the receptor. EV: empty vector as control. (D) Scheme of the constructs of *r*L1 and *h*B3 used in these studies and positions and probability (score, values higher than 0.5 are considered to indicate likely O-glycosylation sites) of O-glycosylation sites predicted by the algorithm of NetOGlyc-4.0 [2].

The theoretical molecular masses based on the construct sequences are 163.2 kDa (177.2kDa with 7 predicted N-glycosylation sites assuming an average mass of 2 kDa) for the full-length rat L1 construct used in this study, 92.8 kDa (106.8 kDa with 7 predicted N-glycans) for the *r*L1 NTF, 172.7 kDa (194.7 kDa with 11 predicted N-glycosylation sites) for the full-length *h*B3 construct and 98.0 kDa for the *h*B3 NTF (118 kDa with 10 predicted N-glycans). Higher masses likely result from O-glycosylation.

We interpret the observations in these WBs as follows. The upper bands in panel A (closed black triangle) correspond to properly folded protein with O- and N-glycosylation. This protein is principally prone to GPS autoproteolysis. The bands marked by the open black triangle correspond to improperly folded protein without O-glycosylation (see below), that is stuck in the ER. Due to the improper fold, the GPS is not active, independent of the GPS sequence. The Δ variant, which lacks a residue at position 836, cannot adopt a proper fold as the *Stachel* sequence is out of register and subdomain B cannot fold. Therefor only unfolded protein, but no cleaved or uncleaved properly folded protein is observed. The other GPS mutations result in proteins with correctly folded GAIN domains, that undergo GPS cleavage to different amounts. WT, F803W, and F803Y are fully cleaved at the GPS, whereas F803A and F803L result in partial cleavage. H836S results in no GPS cleavage. Two bands are observed for the NTF, which result from differentially glycosylated proteins. The analysis in panel C demonstrates, that the bands of likely misfolded protein (open black triangle) correspond to N-glycosylated protein lacking O-glycosylation as treatment with PNGase F ("P"), which removes almost all N-glycans, results in the same shift as treatment with Deglycosylation mix II ("A"), a mixture of enzymes to remove all N-linked and simple O-linked glycans[1]. The bands marked with a closed red triangle contain N- and O-glycosylation, whereas the bands marked with an open triangle contain only N-glycosylated protein. The thickness of the lower band(s) in experiment rL1-A in panel C and its height relative to the lower band of rL1-P indicates that some of the O-glycosylation of the bands marked with the open red triangle has not been removed. Likewise, the two NTF bands of *h*B3 in the experiments in panel B may correspond to protein differing in O-glycosylation. The mutations of residue F803 in *r*L1 (panel A) which reduce GPS cleavage activity also reduce the fraction of protein lacking O-glycosylation. The reason for this behavior is currently not clear.

The deglycosylation enzymes used in experiments "P" and "A" differ also in the activity to remove N-glycans. Whereas PNGase F ("P") cannot remove N-glycans that have an α1-3-linkage to a fucose at the first GlcNAc residue, deglycosylation mix II ("A") would remove such glycans. The *r*L1 construct used for this study contains nine predicted N-glycosylation sites (score > 0.5 using prediction server NetNGlyc-1.0[3]). Assuming an average mass of ~2 kDa per predicted glycosylation site in HEK293T cell expression, this may result in ~18 kDa molecular mass resulting from N-glycosylation. Since deglycosylation with PNGase F results in a mass shift of ~17 kDa and application of the Deglycosylation mix II in a mass shift of ~34 kDa, it appears unlikely that the additional mass shift observed for the deglycosylation mix has large contributions from α1-3-fucosylated GlcNAc residues at the first glycan position.

[1]https://international.neb.com/products/p6044-protein-deglycosylation-mix-ii#Product%20Information

**Table S5: Interfaces between protein chains identified by the PISA and EPPIC servers within the crystal lattice of *h*B2-HG**. The values are derived from the PISA server, values in brackets refer to the results of the EPPIC server. The table lists the number of residues ($N_{RES}$) per monomer that are involved in the interface and how the monomers are symmetry-related to one another. Interfaces are further quantified by their surface area, the free energy (ΔG) gained by formation of the interface and the number of hydrogen bonds ($N_{HB}$), salt bridges ($N_{SB}$) and disulfide bonds ($N_{DS}$) across the interface.

| # | $N_{RES}$(A) | $N_{RES}$(B) | Symmetry relation | Interface area [Å] | ΔG [kcal/mol] | $N_{HB}$ | $N_{SB}$ | $N_{DS}$ |
|---|---|---|---|---|---|---|---|---|
| **1** | 22 (22) | 22 (22) | -x-1, -y, z | 668.6 (669.5) | -2.8 | 0 | 0 | 0 |
| **2** | 17 (17) | 15 (15) | x-1/2, -y+1/2, -z | 496.7 (499.6) | -0.7 | 3 | 9 | 0 |
| **3** | 9 (9) | 9 (9) | -x, -y, z | 444.6 (444.6) | -2.5 | 0 | 6 | 0 |
| **4** | 10 (10) | 14 (13) | -x-1/2, y-1/2, -z-1 | 372.7 (372.3) | 0.5 | 0 | 3 | 0 |
| **5** | 9 (9) | 9 (10) | -x-1/2, y-1/2, -z | 272.3 (271.8) | -3.7 | 0 | 0 | 0 |
| **6** | 4 (4) | 2 (2) | -x-1, -y, z-1 | 51.7 (53.2) | -0.2 | 0 | 0 | 0 |

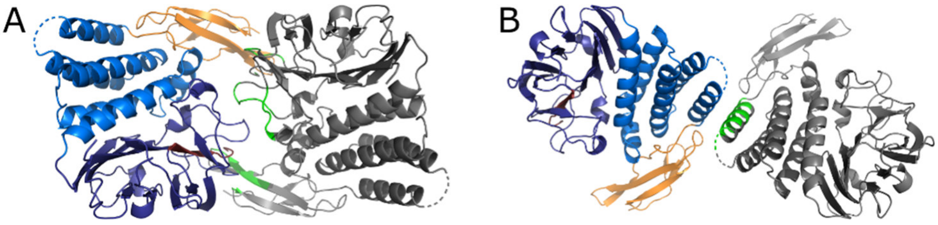


**Figure S22: Fold of the two putative dimer structures with a closed $C_2$ point symmetry observed in the crystal.** (A) Dimer #1 and (B) dimer #3 according to the numbering in Table S6. One protomer is shown with the usual color scheme (HormR domain orange, $GAIN_A$ bright blue, $GAIN_B$ dark blue, *Stachel* dark red), while the other protomer is shown in grey, with residues involved in the interface highlighted in green.

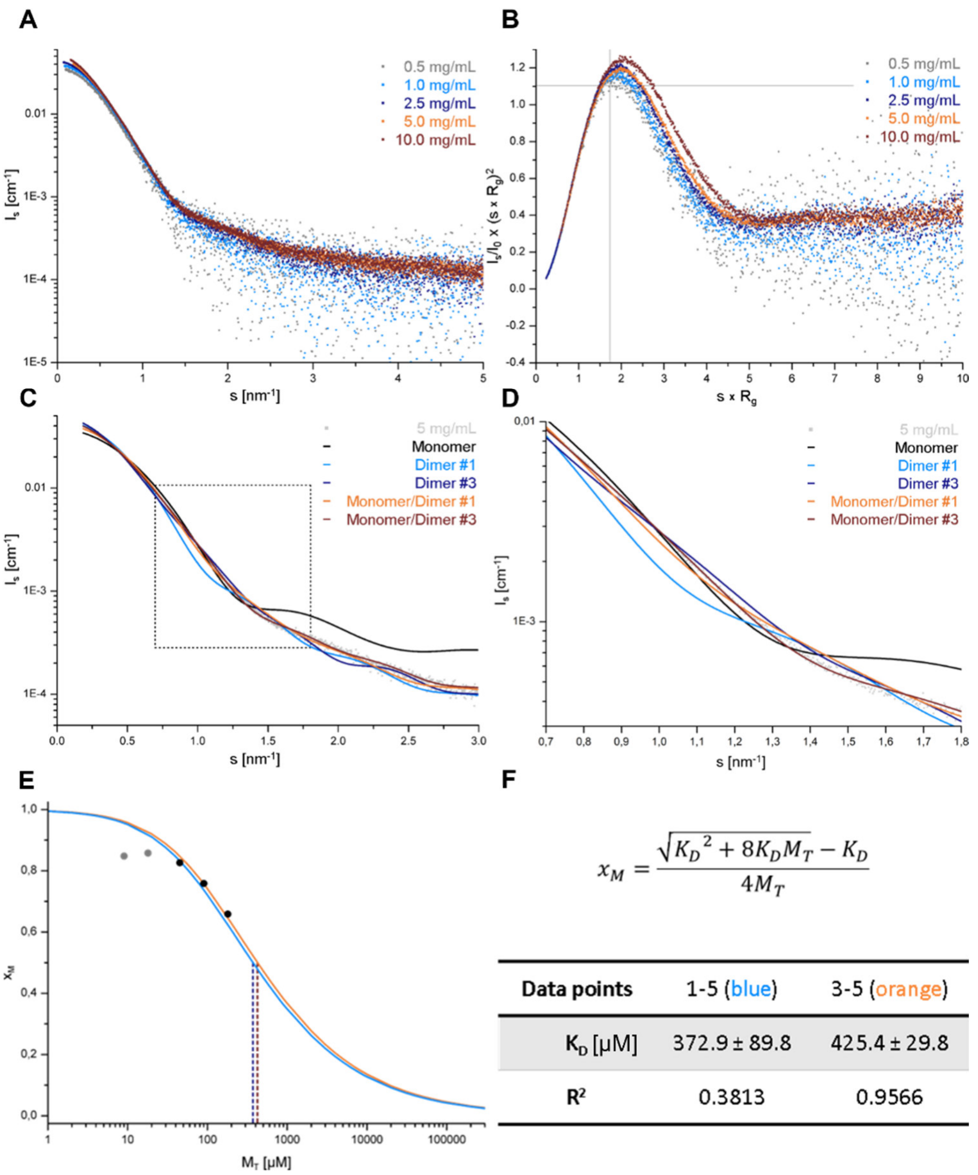


$$x_M = \frac{\sqrt{K_D{}^2 + 8K_DM_T} - K_D}{4M_T}$$

| Data points | 1-5 (blue) | 3-5 (orange) |
|---|---|---|
| $K_D$ [µM] | 372.9 ± 89.8 | 425.4 ± 29.8 |
| $R^2$ | 0.3813 | 0.9566 |

**Figure S23: SAXS data for a dilution series of *h*B2-HG between 0.5 mg/ml and 10 mg/ml.** (A) The logarithmic plot of the absolute intensity against the wave vector s exhibits a change in the scattering profile at low wave vectors with increasing concentration, indicating some form of aggregation or oligomerization. (B) Normalized Kratky plot. The peak maximum shifts to higher concentrations indicating a more elongated particle structure. The grey cross marks the theoretical maximum of $s \cdot R_g = \sqrt{3}$ for a solid sphere. Given the fairly globular fold for the monomer, evident from the crystal structure, this finding is in agreement with oligomerization to an elongated dimeric structure. (C) Fitting of ensemble scattering curves predicted by MultiFoXS to the logarithmic intensity plot at 5 mg/ml suggests that the behavior in solution is best modelled by the presence of monomers and dimer #3. (D) The differentiation between dimer #1 and dimer #3 is mainly due to the differences at s between 1.0 and 1.5 $nm^{-1}$. (E) Estimation of the $K_D$-value for the monomer/dimer equilibrium. The molar fraction of the monomer ($x_M$) as determined from the volume fraction of the SAXS fits is plotted against the total protein concentration ($M_T$). The data was fitted using equation specified in (F). The blue fit curve was obtained using all data points, while for the orange fit only the data points at three highest protein concentrations were used as the data points at low concentrations deviate from the theoretical curve shape corresponding to the law of mass action.

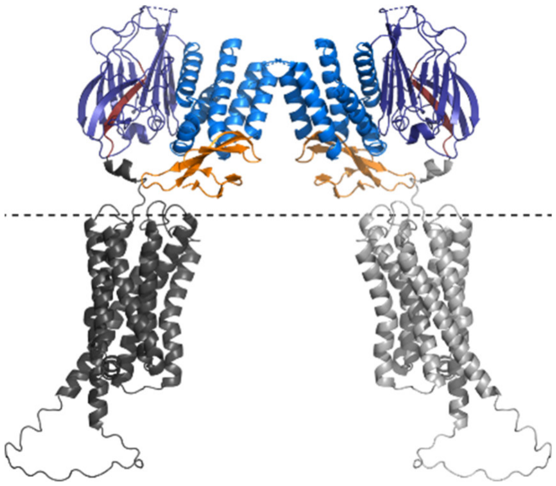

**Figure S24:** Tentative model of the *h*B2-HG dimer complemented with an AlphaFold model of the 7TM domain. The relative positioning of the 7TM region based on an alignment of the full-length AlphaFold *h*B2 model to the crystal structure of the *h*B2-HG dimer. The dashed line indicates the position of the cell membrane. The AlphaFold model of *h*B2 contains a relatively high predicted aligned error (PAE) between the position of the GAIN and the 7TM domain, i.e. the relative positioning of these two domains has low confidence. Thus, the 7TM domain is included in this figure to illustrate the relative size of the domains and the length of the linker region between the *Stachel* sequence and the 7 TM region in the inactive receptor (*Stachel* bound to GAIN domain), while the exact positioning of the 7TM domain is unclear. If the dimer structure observed for *h*B2-HG exists also for the full-length receptor, the HormR domains would be positioned close to the cell membrane.

**Table S6: Comparison of the GPS of GAIN domains to other proteins with *cis*-autoprocessing activity**

| protein | sequence | base | oxoanion hole | protonation amide NH | strain at or near scissile peptide bond | inactivation, pdb id | ref |
|---|---|---|---|---|---|---|---|
| NtN hydrolases: glycosylasparaginase | $^{151}$D↓TI | D151 | T170-OH | D151 suggested*1 | on scissile peptide bond and/or penultimate peptide bond | T152C, 3ljq | 4,5 |
| pantetheine hydrolase ThnT | $^{281}$N↓TT | no | N217-$N_{\delta}H_2$, F144-NH | no | two conformations of main chain in T282C variant. Strained φ,ψ-angles of C282 in state A. Strained ω angle in state B. | T282C, 3s3u | 6 |
| Nup98 | $^{862}$HF↓S | H862 | K791-$NH_3^+$, N799-$NH_2$ | H862?, not discussed in ref.[7] | Strained scissile *cis*-peptide bond | S864A, 2q5x | 7 |
| SEA domains: *h*Muc1 SEA domain | $^{1088}$G↓SVVV | no | no | no | Substantial strain of 7 kcal/mol, scissile bond also strained | modeled | 8,9 |
| Inteins: GyrA | $^{0}$T↓CI*2 | no | T72-OH (N74-$NH_2$) | H75 | Scissile bond is *cis*-peptide, no strain, but *cis*-peptide has ~5 kcal/mol higher energy | $^{0}$A↓SI, 1am2 | 10 |
| Hedgehog proteins: Drosophila hedgehog | $^{256}$HG↓CF | no | T326-OH | H329 | no uncleaved structure available | cleaved: 1at0 | 11 |
| Pyruvoyl enzymes: S-Adenosylmethionine decarboxylase | $^{66}$SE↓SS | no*3 | no, C82-SH? | H243 | no | S68A, 1msv | 12 |

*1D151 was suggested to protonate the leaving amide nitrogen, but it is unclear if a direct proton transfer is possible; *2T is the last residue of the extein; *3the authors suggest that the proton of the serine nucleophile si directly transferred to the carbonyl oxygen of the scissile peptide bond

**Table S7: Primers used for amplification of parts of the BAI2 gene for subcloning into pHLsec and PiggyBac vectors**

| Primer | Sequence | Restriction site |
|---|---|---|
| *h*B2-ECR_33-X | 5'-ACTGAGAGCTCTTCGATCCTGCTCCTTCTGC-3' | *Sac*I |
| *h*B2-THG_296-X | 5'-ATATAGAGCTCAGCGCCGACGAGCCTGGA-3' | *Sac*I |
| *h*B2-HG_528-X | 5'-ATTATGAGCTCCCTGCCTTCCACGAGATGTG-3' | *Sac*I |
| *h*B2_X-921 | 5'-ATATAGGTACCAGGAGGTTGAGCCAGCACG-3' | *Kpn*I |
| *h*B2-C_X-295 | 5'-AGTCAGGTACCTCTAGGCCACTGGGTTTT-3' | *Kpn*I |

**Table S8: Transfection conditions used for all transient expression tests**. PEI: polyethyleneimine, VPA: valproic acid

| **Condition #** | **1** | **2** | **3** | **4** | **5** | **6** |
|---|---|---|---|---|---|---|
| Target DNA/ng | 500 | 500 | 500 | 500 | 500 | 500 |
| pAdVantage DNA/ng | – | – | 100 | 100 | 500 | 500 |
| PEI/ng | 750 | 750 | 900 | 900 | 1500 | 1500 |
| VPA/µM in final volume | – | 4 | – | 4 | – | 4 |

**Table S9. Primers used for construction of *r*L1, *h*B2 and *h*B3 mutants for receptor autoproteolysis assays**

| **Primer** | **Sequence** |
|---|---|
| ac_1F | 5'-TAATACGACTCACTATAGGG-3' |
| ac_3F | 5'-GCTTATGTGATCATCTCTCTACCTTCGCC-3' |
| ac_4R | 5'-GGCGAAGGTAGAGAGATGATCACATAAGC-3' |
| ac_5R | 5'-CCCCTCGAGCCCTCAAACTTCTG-3' |
| ac_21R | 5'-GCTCTAGCATTTAGGTGACAC-3' |
| ac_102F | 5'-AACTGCTCCTGGAACTACTCA-3' |
| ac_103R | 5'-TGAGTAGTTCCAGGAGCAGTT-3' |
| ac_104F | 5'-AACTGCTCCGCCTGGAACTACTCA-3' |
| ac_105R | 5'-TGAGTAGTTCCAGGCGGAGCAGTT-3' |
| ac_106F | 5'-AACTGCTCCTTATGGAACTACTCA-3' |
| ac_107R | TGAGTAGTTCCATAAGGAGCAGTT-3' |
| ac_110F | 5'-AACTGCTCCTGGTGGAACTAC-3' |
| ac_111R | 5'-GTAGTTCCACCAGGAGCAGTT-3' |
| ac_128F | 5'-CTTCAATGCAAACTGCTCCTACTGGAACTACTCAGAGCGC-3' |
| ac_129R | 5'-GCGCTCTGAGTAGTTCCAGTAGGAGCAGTTTGCATTGAAG-3' |
| ac_168F | 5'-CACCTGGCTACCTTTGCCGTGCTG-3' |
| ac_169R | 5'-AAAGGTAGCCAGGTGCTGACACTGGCAT-3' |
| ac_170F | 5'-CAGTGTCAGGCCCTGTCTACC-3' |
| ac_171R | 5'-GGTAGACAGGGCCTGACACTG-3' |
| ac_172F | 5'-CACTGCGCCTTCTGGGATTAC-3' |
| ac_173R | 5'-GTAATCCCAGAAGGCGCAGTG-3' |
